# Dissecting the phase separation and oligomerization activities of the carboxysome positioning protein McdB

**DOI:** 10.1101/2022.04.28.489914

**Authors:** Joseph L. Basalla, Claudia A. Mak, Jordan Byrne, Maria Ghalmi, Y Hoang, Anthony G. Vecchiarelli

## Abstract

Across bacteria, protein-based organelles called bacterial microcompartments (BMCs) encapsulate key enzymes to regulate their activities. The model BMC is the carboxysome that encapsulates enzymes for CO_2_ fixation to increase efficiency and is found in many autotrophic bacteria, such as cyanobacteria. Despite their importance in the global carbon cycle, little is known about how carboxysomes are spatially regulated. We recently identified the two-factor system required for the maintenance of carboxysome distribution (McdAB). McdA drives the equal spacing of carboxysomes via interactions with McdB, which associates with carboxysomes. McdA is a ParA/MinD ATPase, a protein family well-studied in positioning diverse cellular structures in bacteria. However, the adaptor proteins like McdB that connect these ATPases to their cargos are extremely diverse. In fact, McdB represents a completely unstudied class of proteins. Despite the diversity, many adaptor proteins undergo phase separation, but functional roles remain unclear. Here, we define the domain architecture of McdB from the model cyanobacterium *Synechococcus elongatus* PCC 7942, and dissect its mode of biomolecular condensate formation. We identify an N-terminal intrinsically disordered region (IDR) that modulates condensate solubility, a central coiled-coil dimerizing domain that drives condensate formation, and a C-terminal domain that trimerizes McdB dimers and provides increased valency for condensate formation. We then identify critical basic residues in the IDR, which we mutate to fine-tune condensate solubility. Finally, we find that a condensate-defective mutant of McdB has altered association with carboxysomes and influences carboxysome enzyme content. The results have broad implications for understanding spatial organization of BMCs and the molecular grammar of protein condensates.

## INTRODUCTION

Compartmentalization is a fundamental feature by which cells regulate metabolism. Although bacteria lack extensive lipid-membrane systems, recent reports have shown that proteinaceous <u>b</u>acterial <u>m</u>icro<u>c</u>ompartments (BMCs) are a widespread strategy for compartmentalization in bacteria (1, 2). Briefly, BMCs are nanoscale reaction centers where key enzymes are encapsulated within a selectively-permeable protein shell. The best studied BMC is the carboxysome, found within cyanobacteria and other autotrophic bacteria (1, 3). Carboxysomes encapsulate the enzyme ribulose-1,5-bisphosphate carboxylase/oxygenase (Rubisco) with its substrate CO_2_ to significantly increase the efficiency of carbon fixation. Carboxysomes serve as a paradigm for understanding BMC homeostasis, including assembly, maintenance, permeability, and spatial regulation (1, 2, 3). Furthermore, to engineer efficient carbon-fixing organisms, efforts to express functional carboxysomes in heterologous hosts are ongoing (4, 5).

An important aspect of BMC homeostasis is spatial regulation (6). We recently identified the two-component system responsible for spatially regulating carboxysomes, which we named the <u>m</u>aintenance of <u>c</u>arboxysome <u>d</u>istribution (McdAB) system (7, 8, 9). McdA and McdB function to prevent carboxysome aggregation, thereby ensuring optimal function and equal inheritance upon cell division (7, 9). Briefly, McdA is an ATPase that forms dynamic gradients on the nucleoid in response to an adaptor protein, McdB, which associates with carboxysomes (7, 9). The interplay between McdA gradients on the nucleoid and McdB-bound carboxysomes result in the equal spacing of carboxysomes down the cell length of rod-shaped bacteria. This mode of spatial regulation by McdA is typical for the widespread and well-studied ParA/MinD family of positioning ATPases, of which McdA is a member. ParA/MinD ATPases spatially organize an array of genetic- and protein-based cargos in the cell, including plasmids, chromosomes, the divisome, flagella, and other mesoscale complexes (10, 11). While ParA/MinD ATPases are highly similar in sequence and structure, the partner proteins that act as adaptors and link the ATPases to their respective cargo are highly diverse, largely due to partners providing cargo specificity. Indeed, McdB represents an entirely new class of adaptor proteins, and it is therefore unknown how McdB interacts with itself, McdA, and carboxysomes to confer specificity, or how these interactions are regulated. Bioinformatic analyses show that McdAB systems also exist for several other BMC types (9). Therefore, an understanding of the biochemical properties of McdB, and how these properties influence its behavior *in vivo,* are important next steps to advancing our knowledge on the spatial regulation of carboxysomes and BMCs in general.

From our initial studies in the model cyanobacterium *<u>S</u>ynechococcus <u>e</u>longatus* PCC <u>7942</u> (*Se7942*), we found that McdB self-associates *in vitro* to form both a stable hexamer (9) and liquid-like condensates (8). However, the domain architecture of McdB and the regions required for its oligomerization and condensate formation are unknown. Our understanding of how proteins form biomolecular condensates has rapidly developed over the past decade. Briefly, biomolecular condensates are the result of molecules having demixed out of solution to form a dense, solvent-poor phase that exists in equilibrium with the soluble phase (14, 15, 16). This process occurs under a specific set of conditions where protein-protein interactions are more favorable than protein-solvent interactions (15). A biochemical understanding of how proteins form condensates *in vitro* has led to a deeper understanding of how this process facilitates subcellular organization in both eukaryotic and prokaryotic cells (17, 18). Furthermore, characterizing the underlying chemistries for diverse biomolecular condensates has led to the development of these condensates as synthetic tools to engineer cytoplasmic organization (19,20, 21, 22). Thus, a major focus of this report is to characterize the biochemistry of McdB, including its condensate formation, and link these properties to the spatial regulation of carboxysomes *in vivo*.

Here we define a domain architecture of *Se7942* McdB, identify the domains contributing to oligomerization and condensate formation, and discover a potential interplay between these two modes of self-association. We then create a series of point mutations that allow us to fine-tune the solubility of McdB condensates both *in vitro* and *in vivo* without affecting McdB structure or oligomerization. Finally, we use this mutation set to identify *in vivo* phenotypes that relate specifically to the ability of McdB to form condensates and associate with carboxysomes. The findings have implications for the use of carboxysomes in synthetic biology approaches, designing biomolecular condensates, and general BMC biology.

## RESULTS

### Structural predictions generate a low confidence α-helical model for Se7942 McdB

We first set out to determine the *Se7942* McdB crystal structure. However, McdB displayed robust phase separation across a range of buffer conditions, making crystal trials thus far unsuccessful (Fig. S1). We next turned to I-TASSER (Iterative Threading ASSEmbly Refinement) (24, 25) to generate structural models, which predicted the McdB secondary structure to be predominantly α-helical but with a disordered N-terminus (Fig. S2A). I-TASSER also generates full-length atomic models of the target sequence that are consistent with its secondary structure predictions via multiple sequence alignments using top matches from the protein databank (PDB). The top three models were once again almost entirely α-helical, with the top model also showing a disordered N-terminus (Fig. S2B). But ultimately, the top 10 PDB matches identified by I-TASSER aligned poorly with McdB, with each alignment showing low sequence identity (< 20% on average) and low-quality scores (Z-scores < 1 on average) (Fig. S2C). As a result, the top three final models generated by I-TASSER all have poor confidence scores (Fig. S2B). These findings are not surprising in context with our previous bioinformatic analyses showing that cyanobacterial McdBs are highly dissimilar to other characterized proteins at the sequence level, potentially related to the high disorder content of McdBs (8, 26). Together, these data provide low-confidence structural predictions for *Se7942* McdB. We therefore set out to validate these predictions with empirical approaches.

### Defining a tripartite domain architecture for Se7942 McdB

We used circular dichroism (CD) to characterize the secondary structure of *Se7942* McdB. The spectrum showed a characteristic α-helical signature that remained stable even after incubation at 80°C (Fig. 1A), indicative of a stabilized coiled-coil (27). This is consistent with the helical predictions from I-TASSER, and with our previous bioinformatics data that predicted coiled-coil domains to be conserved across all cyanobacterial McdB homologs (8).

**Figure 1:**
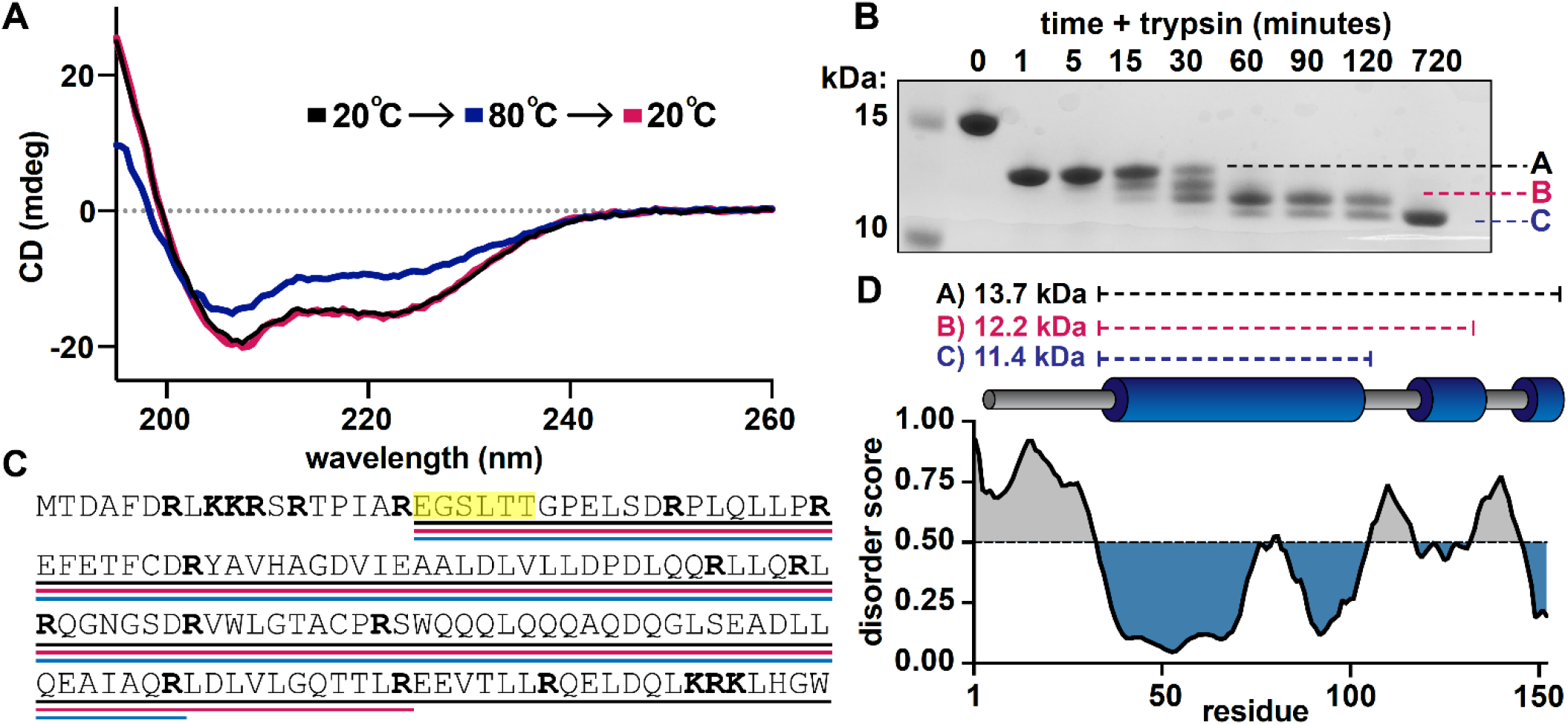
Defining a domain architecture of *Se7942* McdB. (**A**) Circular Dichroism (CD) spectra of McdB at 20°C (black), 80°C (blue), and then returned to 20°C (magenta). Spectra show α-helical structure resilient to heat denaturation. (**B**) SDS-PAGE analysis of trypsin-digested McdB sampled over digestion time. Bands labeled A, B, and C were isolated and N-terminally sequenced. (**C**) Amino acid sequence of McdB with basic residues (Lys-K and Arg-R) in bold. Regions corresponding to bands A, B, and C from panel B are underlined in black, magenta, and blue, respectively. Amino acids determined through N-terminal sequencing of bands A, B, and C are highlighted yellow. (**D**) Structural model of *Se7942* McdB. Regions corresponding to bands A, B, and C are indicated with predicted MWs (*top*). Predicted secondary structure of McdB (*middle*) aligned with a <u>P</u>redictor <u>o</u>f <u>N</u>atural <u>D</u>isorder <u>R</u>egions (PONDR) plot using the VLXT algorithm (*bottom*) with disordered regions colored grey and predicted α-helical domains in blue.

We next sought to empirically identify folded domains using limited proteolysis (28, 29, 30) (Fig. 1B). Trypsin cuts at arginines and lysines, which are frequent throughout *Se7942* McdB – the largest fragment between any two basic residues is ∼3 kDa (Fig. 1C). Therefore, any stably folded regions that are protected from trypsin would be resolved via this approach. The digestion yielded three major bands of varying stabilities, which we labeled A, B, and C (Fig. 1B). Band C was most stable, representing an ∼11 kDa fragment that remained undigested for 12 hours. This strong protection against trypsin is consistent with our CD data, which showed high resilience to heat denaturation (see Fig. 1A).

We next subjected bands A, B, and C to N-terminal sequencing to determine the location of these stably folded regions in McdB. All three bands had the same N-terminal sequence starting at E19 (Fig. 1C). Therefore, the first 18 amino acids at the N-terminus were digested within the first minute to produce band A, and further digestion progressed slowly from the C-terminus to produce bands B and C (Fig. 1B, C). By combining the N-terminal sequencing results (Fig. 1C), the molecular weights of the three protected regions (Fig. 1B), the locations of all arginines and lysines (Fig. 1C), and the predicted disorder via PONDR VLXT (31) (Fig. 1D), we developed a model for the domain architecture of *Se7942* McdB that was consistent with I-TASSER predictions (Fig. 1D).

### Se7942 McdB forms a trimer-of-dimers hexamer

From our structural model, we defined three major domains of *Se7942* McdB: (1) an intrinsically <u>d</u>isordered <u>r</u>egion (IDR) at the N-terminus, (2) a highly stable central coiled-coil (CC), and (3) a <u>C</u>-terminal <u>d</u>omain (CTD) with two short helical regions. We used this model to design a series of truncation mutants, including each of these domains alone as well as the CC domain with either the N-terminal IDR or the CTD (Fig. 2A). CD spectra of these truncations showed that the N-terminus was indeed disordered on its own, and both the CC domain and CTD maintained α-helical signatures (Fig. 2B, Fig. S3A).

**Figure 2:**
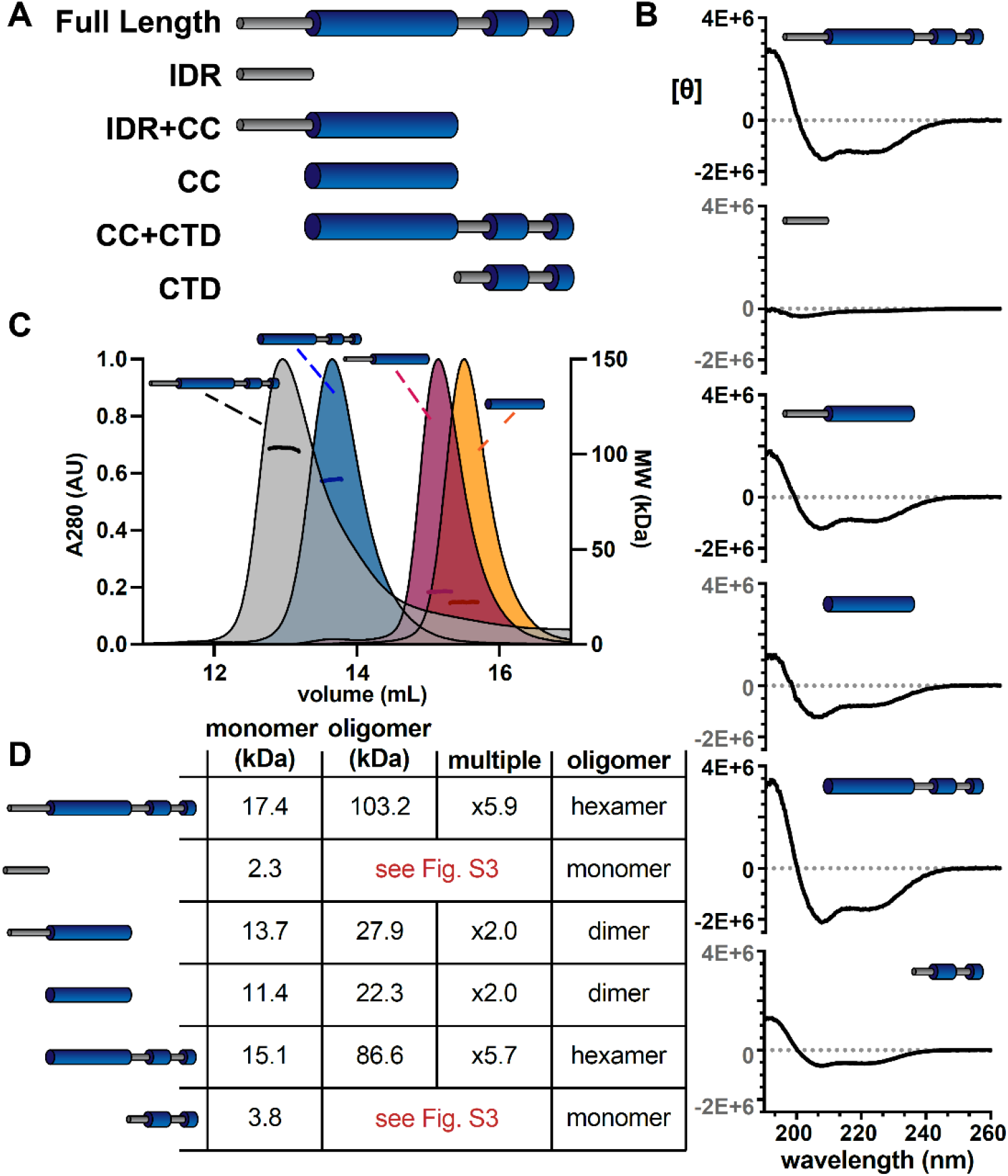
The α-helical domains of McdB form a trimer-of-dimers hexamer. (**A**) Illustration of McdB truncations generated based on the predicted domain structure. (**B**) CD spectra normalized by MW for the indicated McdB truncations. Spectra show α-helical content for all truncations, except for the disordered N-terminal fragment. (**C**) <u>S</u>ize <u>e</u>xclusion <u>c</u>hromatography coupled to <u>m</u>ulti-<u>a</u>ngle light <u>s</u>cattering (SEC-MALS) for full-length McdB and truncation mutants that showed oligomerization activity **(see Fig. S3)**. (**D**) Summary of the SEC-MALS data from (C).

Having previously shown that full-length *Se7942* McdB forms a hexamer in solution (9), we used these truncations to determine which domains contributed to oligomerization. We first ran size-exclusion chromatography (SEC) with each McdB truncation, and used the full-length protein as a reference for where the hexamer elutes. Although the molecular weight of each monomeric truncation is within ∼5 kDa of one another (Fig. S3B), we found that only the CC+CTD construct eluted at a volume similar to the full-length hexamer (Fig. S3C). Furthermore, the CC domain alone or with the IDR, appeared to elute between the expected monomer and hexamer peaks, suggesting that the CC domain with or without the IDR forms an oligomeric species that is smaller than a hexamer.

To further resolve the oligomeric states of these McdB truncations, we performed size-exclusion chromatography coupled to multi-angled light scattering (SEC-MALS) on each of the constructs that appeared to oligomerize during SEC (Fig. 2C). We found that the CC+CTD truncation was indeed hexameric, while the CC domain, with or without the IDR, formed a dimer (Fig. 2D). These data suggest that the CC domain of McdB contains a dimerization interface, while the CTD subsequently allows for trimerization of dimers. Although we were unable to generate an atomic level structure of McdB nor determine the orientation of monomers within the hexamer, we have identified two key oligomerizing domains and conclude that full-length *Se7942* McdB forms a hexamer as a trimer-of-dimers.

### Se7942 McdB forms condensates via pH-dependent phase separation coupled to percolation

Recent reports on the formation of protein condensates have begun to unveil an interplay between phase separation and network formation or “percolation” (32, 16). These reports have shown that, instead of forming strictly through liquid-liquid phase separation, many proteins undergo <u>p</u>hase <u>s</u>eparation <u>c</u>oupled to <u>p</u>ercolation (PSCP) to form condensates (32, 16). The definitions of phase separation, percolation, and PSCP are rigorous and nuanced. But broadly speaking, phase separation can often involve incompatibilities in solubility to drive transitions in density, while percolation concerns multivalent interactions to form dense networks (16). For protein systems undergoing PSCP, different regions of the protein can facilitate solubility than the regions that facilitate multivalent networking (16). We therefore set out to identify if full-length McdB showed signs of PSCP to guide our investigation on how the different domains of McdB affect condensate formation and stability.

Evidence for PSCP has recently been shown by studying the time-dependent viscoelastic nature of condensates (33, 34, 16) and the formation of networks at subsaturating concentrations (32, 16). We therefore first used fluorescence recovery <u>a</u>fter <u>p</u>hotobleaching (FRAP) to assess the viscoelastic nature of McdB condensates (15). Newly formed condensates recovered within minutes (Fig. 3A), and readily fused and relaxed into spheres within seconds (Fig. 3B), reminiscent of liquid-like fluid droplets. But after 18 hrs, recovery was significantly slower (Fig. 3A) and these ‘mature’ droplets no longer fused. The time-dependent changes in material properties of McdB condensates is a signature of PSCP (16).

**Figure 3:**
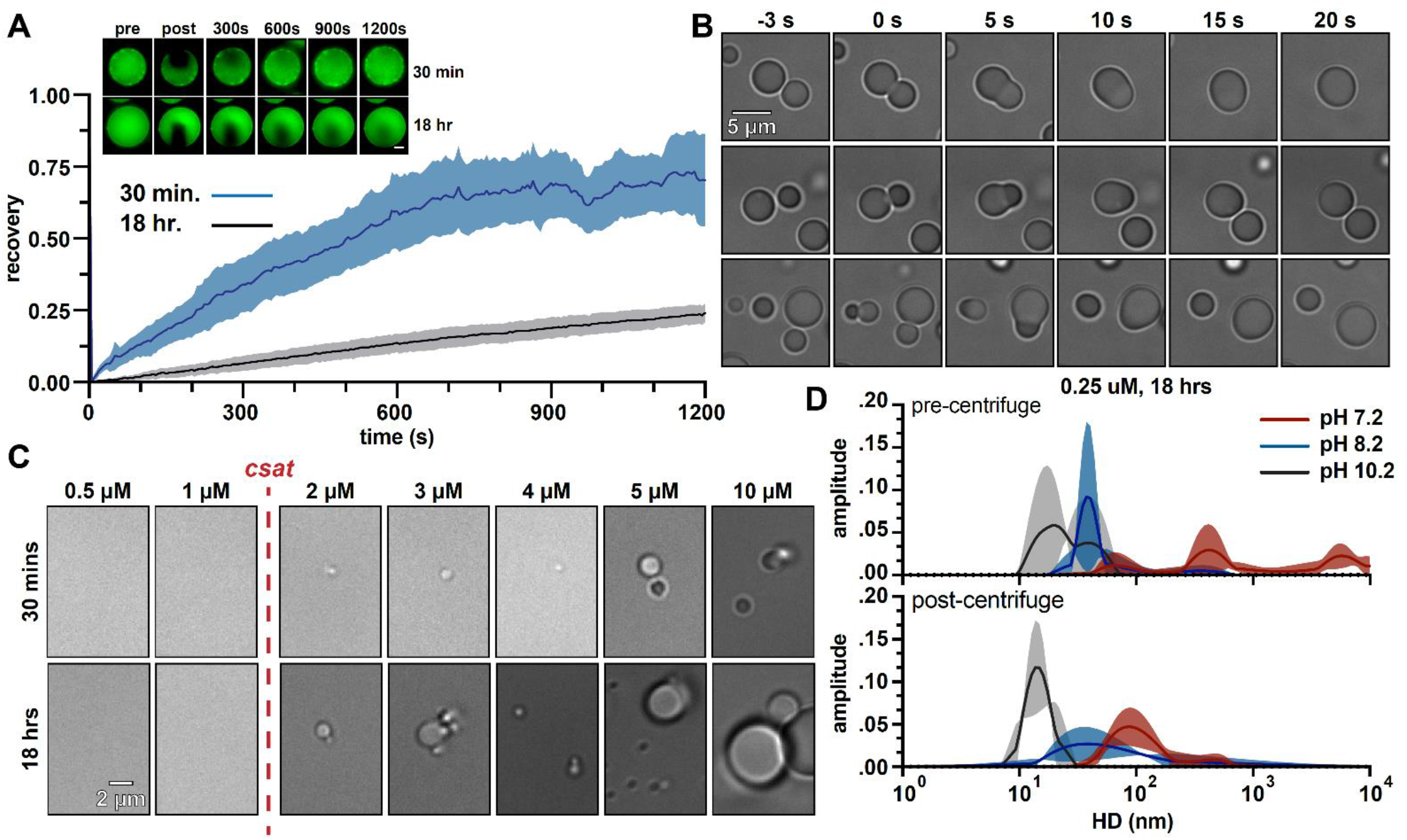
McdB from *Se7942* forms liquid-like condensates via pH dependent phase separation coupled to percolation (PSCP). (**A**) Fluorescence-Recovery After Photobleaching (FRAP) of McdB condensates at the indicated time points. Means and SD from n=8 condensates are shown. Representative fluorescence microscopy images for condensates incubated at 30 mins. or 18 hr. are shown (*inlet, scale bar = 2 µm*) (**B**) Representative DIC microscopy timeseries showing newly formed McdB condensates fusing and relaxing into spheres on the order of seconds. Scale bar applies to all images. (**C**) Representative DIC microscopy images at the indicated protein concentrations. McdB condensates were seen at and above concentrations of 2 µM, suggesting a saturation concentration (c_sat_) between 1 – 2 µM. Scale bar applies to all images. (**D**) Dynamic Light Scattering (DLS) of McdB at a concentration ∼1/10 the c_sat_ determined from (C) and at increasing pH values as indicated. Samples were analyzed both before (*top*) and after (*below*) a 5 min. spin at 20,000 x g. Larger “networks” are seen forming at lower pHs, even below the observed csat.

Next, we determined a saturation concentration (c_sat_) for McdB condensate formation in our standard buffer conditions (100 mM KCl, 20 mM HEPES, pH 7.2). Condensates were observed at or above 2 µM, suggesting a c_sat_ between 1-2 µM (Fig. 3C). We then used <u>d</u>ynamic light <u>s</u>cattering (DLS) to investigate whether McdB formed network-like species at concentrations below c_sat_ as done previously for other proteins (32). At 0.25 µM (∼1/10 the observed c_sat_) in our standard buffer (pH 7.2), McdB displayed a heterogeneous size distribution of defined species spanning 100 – 1000 nm in diameter (Fig. 3D); significantly larger than a monodispersed hexamer (35). Even after high-speed centrifugation, McdB in the supernatant remained mainly as mesoscale clusters on the order of 100 nm, suggesting the formation of McdB clusters at pH 7.2 (Fig. 3D).

Our previous work has shown that McdB condensates are solubilized at higher pH values (8), therefore we set out to determine if McdB clusters remained in solution at higher pH. Interestingly, the clusters were pH-dependent. At pH 8.2, the largest clusters (> 100 nm) were lost, but smaller clusters (∼ 60 nm) remained. At pH 10.2, a single homogeneous species remained at ∼ 10 nm (Fig. 3D), which is consistent with the hydrodynamic diameter of a monodispersed McdB hexamer (35). Together, the data suggest *Se7942* McdB forms condensates via pH-dependent PSCP.

### The CC domain *of* Se7942 McdB is necessary and sufficient for condensate formation

We next used the truncations to determine how each domain of *Se7942* McdB affects condensate formation and stability. Interestingly, no McdB truncations formed condensates under the buffer conditions that sufficed for full-length McdB (Fig. 4A). This finding suggests that no single domain of McdB is sufficient for full-length level condensate formation. Rather, all domains must influence McdB condensates to some extent.

**Figure 4:**
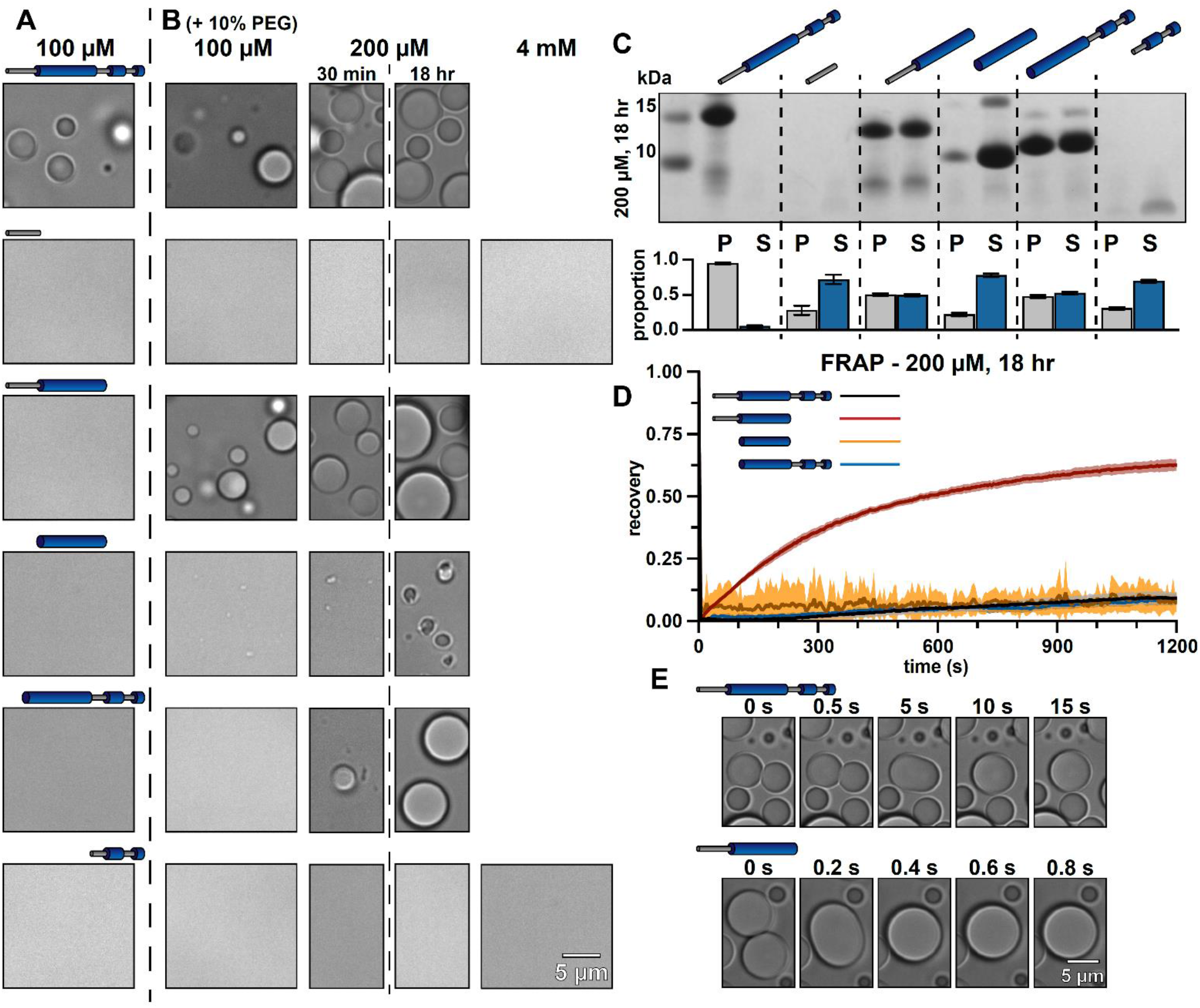
Truncations provide insight into the mechanisms of McdB condensate formation and stabilization. (**A**) Representative DIC microscopy images of full-length and truncation mutants of McdB at 100 µM in 150 mM KCl and 20 mM HEPES pH 7.2. (**B**) As in (A), but with increasing protein concentration as indicated and with the addition of 10% PEG-8000. Scale bar applies to all images. All domains are required for FL level condensate formation (**C**) Condensates at 200 µM after 18 hrs were pelleted (P) and run on an SDS-PAGE gel along with the associated supernatant (S) (*top*). P and S band intensities were then quantified (*bottom*). Mean and SD from 3 replicates are shown. (**D**) FRAP of condensates at the indicated condition reveal an increase in dynamics when the N-term IDR is present without the CTD. (**E**) Condensates containing the N-term IDR fuse orders of magnitude more quickly in the absence of the CTD, suggesting a stabilizing interaction between the two termini.

When we added a crowding agent (10% PEG) to increase the local protein concentration, both the IDR and CTD alone were unable to form condensates, even at concentrations up to 4 mM (Fig. 4B). However, all truncations containing the CC domain formed condensates. In fact, the CC domain alone was necessary and sufficient for forming condensates, albeit at much higher concentrations than full-length McdB. McdB condensates formed and fused similarly in the presence of other crowding agents, showing these activities were not PEG specific (Fig. S4). The data implicate the CC domain as the driver of McdB condensate formation.

### The IDR and CTD domains are modulators of McdB condensate formation

Although the IDR and CTD were not required for the CC domain to form condensates, fusing either of these domains back onto the CC increased condensate formation and size (Fig. 4B). By using centrifugation to quantify the amount of protein in the dense versus light phases (15), we found that the addition of either the IDR or CTD onto the CC domain comparably increased condensate formation (Fig. 4C). However, by performing FRAP on ‘mature’ condensates (incubated for 18 hrs), we found that the IDR+CC condensates recovered much faster than all other constructs, including full-length McdB (Fig. 4D). Moreover, newly formed IDR+CC condensates fused and relaxed into spheres an order of magnitude faster than newly formed full-length McdB condensates (Fig. 4E).

Together, we draw the following conclusions: (*i*) The CC domain is necessary and sufficient for condensate formation, although to a lesser extent than full-length McdB; (*ii*) The IDR increases solvent interactions, thus affecting phase separation of the CC domain, but seemingly not percolation. This is supported by the fact that the IDR+CC construct has increased condensate formation compared to the CC alone. However, IDR+CC condensates do not mature, lacking the change in viscoelasticity seen for full-length McdB (see Fig. 3A); (*iii*) the CTD increases multivalent interactions to support condensate formation via PSCP. This occurs ostensibly by increasing oligomerization, which in turn increases valency (36, 37), and by the CTD itself providing network-forming contacts within condensates; and lastly (*iv*) the fact that full-length McdB, which contains the IDR and CTD, does not show the same fluid-like behavior as the IDR+CC (no CTD), suggests that the CTD may interact with the IDR within condensates formed by full-length McdB. Using this information, we next sought out residues that affect condensate formation, but not McdB structure or its ability to form a hexamer.

### Net charge of the IDR modulates McdB condensate solubility

To determine which types of residues influence condensate formation, we first performed turbidity assays across a range of protein concentrations, salt concentrations, and pH as previously described (38, 15). Over all McdB concentrations, the phase diagrams showed decreased turbidity at higher KCl concentrations (Fig. S5), implicating electrostatic interactions. We also found that turbidity decreased at higher pH (Fig. S5), suggesting that positively charged residues are important in the solubilization of condensates. We used centrifugation to quantify the amount of McdB in the dense-versus light-phases across KCl and pH titrations while keeping McdB concentration constant. Again, we found a clear increase in the soluble fraction and decreases in condensate size and number as both KCl (Fig. 5A) or pH was increased (Fig. 5B). The data reveal a critical role for positively charged residues in McdB condensate stability.

**Figure 5:**
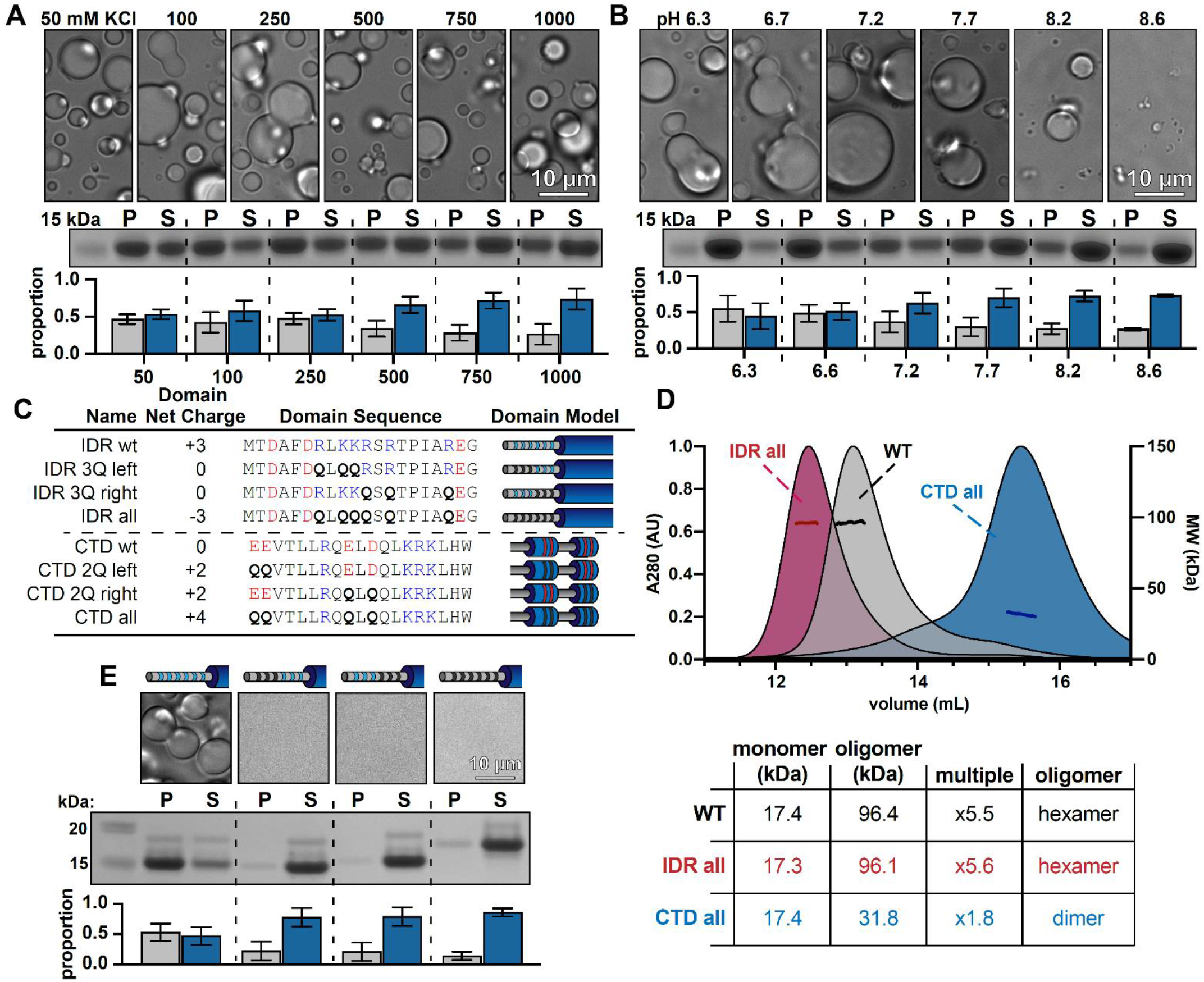
McdB condensates can be solubilized by mutating basic residues in the N-terminal IDR without affecting McdB structure. (**A**) Representative DIC microscopy images of 50 µM McdB in 20 mM HEPES pH 7.2 and increasing KCl concentration (*top*). Scale bar applies to all images. McdB condensates were pelleted (P) and run on an SDS-PAGE gel along with the associated supernatant (S) (*middle*). P and S band intensities were then quantified (*bottom*). Mean and SD from 3 replicates are shown (**B**) As in (A), except salt was held constant at 100 mM KCl and the pH was increased as indicated. (A) and (B) implicate stabilizing basic residues **(see Fig. S6)** (**C**) Table showing the net charge and amino acid sequence of wild-type McdB compared to the glutamine (Q) -substitution mutants in both the N-term IDR and CTD. Acidic and basic residues in the IDR are colored red and blue, respectively. Q-substitutions are bolded. Graphical models of the McdB variants are also provided. (**D**) SEC-MALS of WT McdB compared to the full Q-substitution mutants from both the N-and C-termini. (*Below*) Table summarizing the SEC-MALS data, showing that mutations to the IDR does not affect oligomerization, while mutations to the CTD destabilize the trimer-of-dimers hexamer **(see Fig. S7)**. (**E**) Representative DIC microscopy images for WT and IDR Q-substitution mutants of McdB at 100 µM in 150 mM KCl and 20 mM HEPES pH 7.2 (*top*). Scale bar applies to all images. McdB condensates were pelleted (P) and run on an SDS-PAGE gel along with the associated supernatant (S) (*middle*). P and S band intensities were then quantified (*bottom*). Mean and SD of 3 replicates are shown.

Together with our previous data showing that both the IDR and CTD modulate condensation, we focused on the basic residues within these two domains. By making a series of alanine substitutions (Fig. S6A), we found that removing positive charge in the IDR, but not the CTD, caused a loss of condensates (Fig. S6B). However, we also found that substituting charged residues for a more hydrophobic residue like alanine caused protein aggregation (Fig. S6B) as found for other proteins (39). Therefore, going forward, we transitioned to substituting these charged residues with polar glutamines (Fig. 5C), to specifically affect charge and not hydrophilicity.

Data from this report and our previous study (8) suggest a potential electrostatic interaction between the positively-charged IDR and negatively-charged residues of the CTD. We therefore created another series of substitutions where we changed either positive charge in the IDR or negative charge in the CTD to glutamines. Before assessing condensate formation, we first performed SEC-MALS to verify these mutations had no major impact on McdB structure. Substituting all six basic residues to glutamines in the IDR had no effect on McdB hexamerization (Fig. 5D). On the other hand, only two substitutions in the CTD were enough to partially destabilize the hexamer (Fig. S7), and four substitutions produced mainly McdB dimers (Fig. 5D). As a result, we were unable to parse out the different roles of the CTD in McdB oligomerization versus potential interactions with the IDR involved in condensate formation. Importantly, however, we determined that removing only three positively charged residues in the IDR solubilized McdB condensates (Fig. 5E) without affecting protein structure (Fig. S8) or hexamerization (Fig. 5D).

### McdB condensate formation is tunable through changes in IDR net charge

Substituting only three basic residues in the IDR (net charge -3) completely solubilized McdB condensates under our standard conditions (Fig. 5E). We set out to determine the effect of fewer mutations in the IDR on condensate solubility. Pairs of basic residues in the IDR were substituted with glutamines, leaving an IDR net charge of +1 (Fig. 6A). All +1 mutants still formed condensates, albeit smaller and fewer than that of wildtype McdB (Fig. 6B).

**Figure 6:**
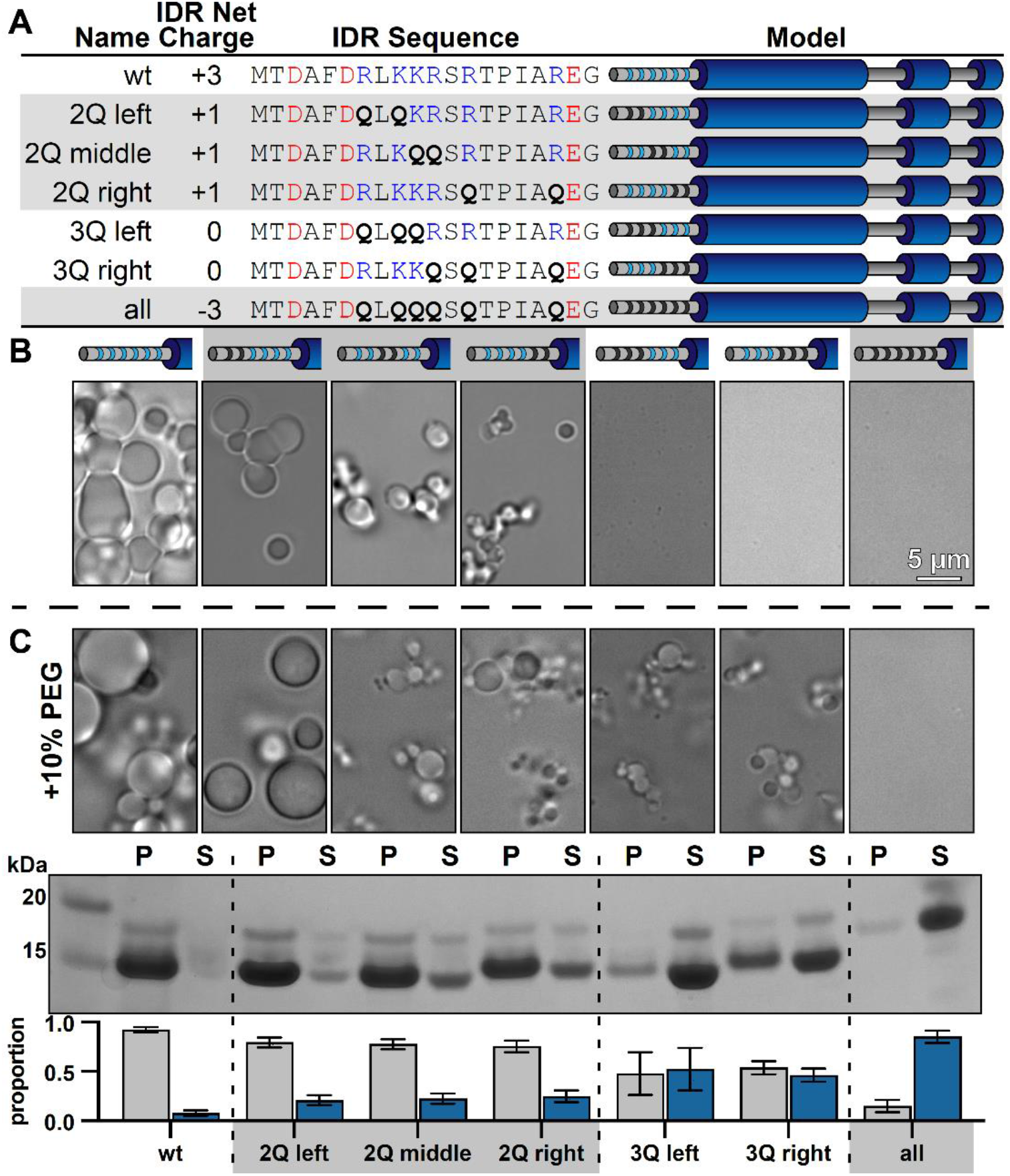
Net charge of the IDR can be used to tune the solubility of McdB condensates. (**A**) Table showing the net charge and N-terminal IDR sequence of wild-type McdB compared to the glutamine (Q)-substitution mutants. Acidic and basic residues in the IDR are colored red and blue, respectively. Q-substitutions are bolded. Graphical models of the McdB variants are also provided where blue stripes represent the six basic residues in the IDR. Black stripes represent the location of the Q-substitutions. (**B**) Representative DIC microscopy images for wild-type and the Q-substitution mutants of McdB at 100 µM in 150 mM KCl and 20 mM HEPES pH 7.2. Scale bar applies to all images. (**C**) As in (B), but with the addition of 10% PEG-8000 (*top*). McdB condensates were pelleted (P) and run on an SDS-PAGE gel along with the associated supernatant (S) (*middle*). P and S band intensities were then quantified (*bottom*). Mean and SD from 3 replicates are shown.

The data suggest McdB condensate formation is tunable though changes to the net charge of the IDR. If correct, the triplet substitution mutants (IDR net charge 0) may still be capable of forming condensates at higher protein concentrations. Indeed, when we added a crowding agent, the net-charge 0 mutants formed condensates (Fig. 6C). Moreover, a gradual increase in the proportion of McdB in the soluble phase was revealed as we incrementally removed positive charge from the IDR. Removing all six positive residues (net charge +3) still completely solubilized McdB even in the presence of a crowder (Fig. 6C). Importantly, McdB mutants with the same IDR net charge, but with different residues substituted, showed similar changes to condensate solubility (Fig. 6C). Substitution position showed slight differences in condensate size, but the overall effect on solubility was the same within each charge grouping. Together, the data show that it is the net charge of the IDR, and not a specific basic residue, that is critical for mediating condensate solubility.

### Net charge of the IDR affects McdB condensation in E. coli

To determine if the IDR can be used to tune McdB solubility in cells, we induced expression of fluorescent fusions of mCherry with both wildtype McdB (McdB[wt]) and the full glutamine-substitution mutant, with an IDR net charge of -3 (“McdB[-3]”) in *E. coli* MG1655. As protein concentration increased, McdB[wt] formed polar foci that coexisted with a dilute cytoplasmic phase (Fig. 7A), similar to the dense and dilute phases of McdB *in vitro*. After 3 hours of expression, nearly 70% of cells with McdB[wt] adopted this two-state regime (Fig. 7B). The foci were indeed driven by McdB, as mCherry alone remained diffuse (Fig. 7A-B). McdB[-3], on the other hand, was considerably more soluble than wild-type, where even after 3 hours of expression <10% of cells contained foci (Fig. 7A-B). The change in solubility was not due to differences in protein levels or due to cleavage of the fluorescent tag (Fig. 7C), but instead represents an increased solubility due to the IDR substitutions. Together the data show that adjustments to the net charge of the IDR can also affect McdB condensate solubility *in vivo*.

**Figure 7:**
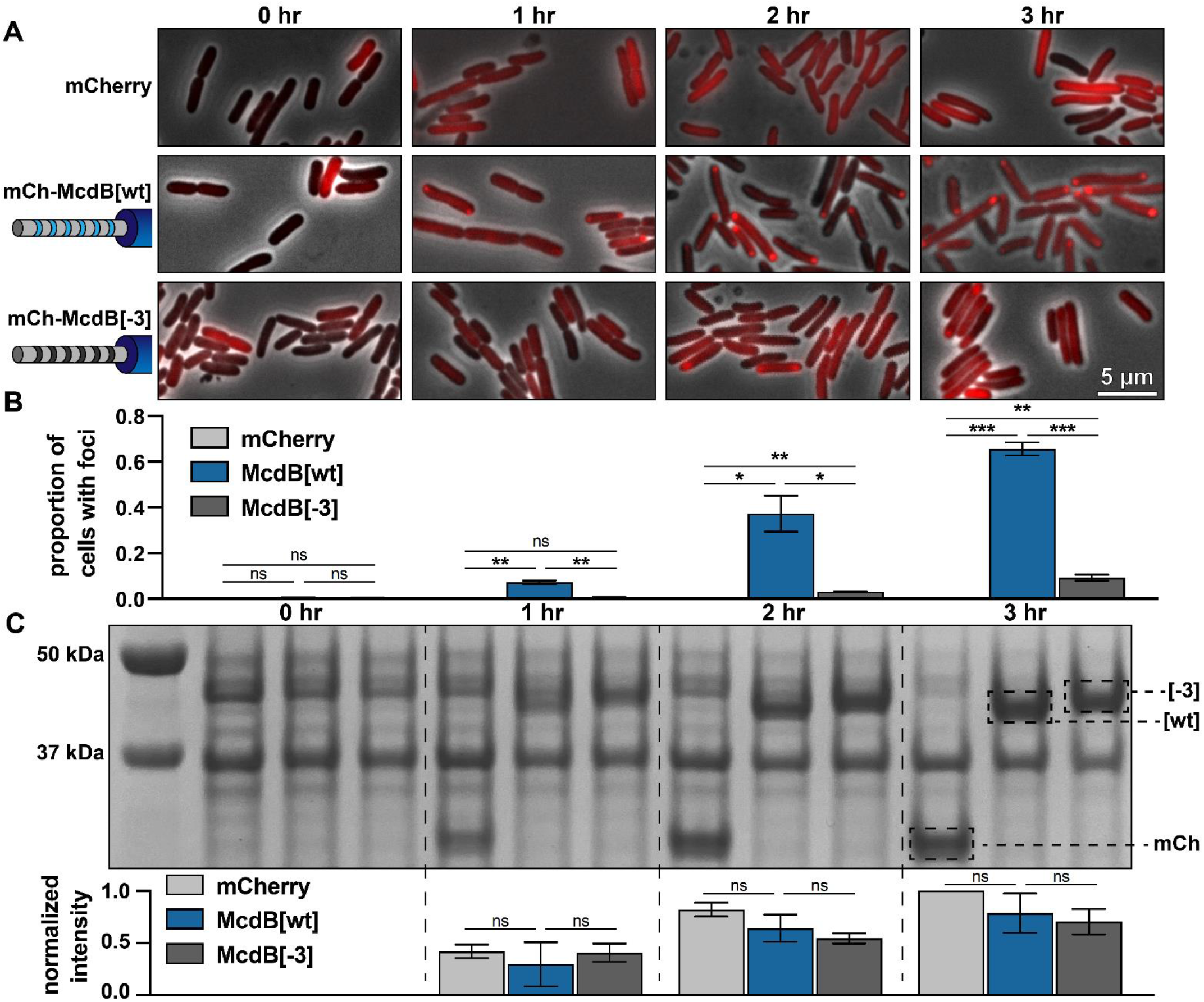
Net charge of the IDR affects McdB solubility in *E. coli*. (**A**) Representative fluorescence microscopy images monitoring the expression of the indicated constructs over time. Scale bar applies to all images. (**B**) Quantification of the proportion of cells containing foci from the images represented in (A). All quantifications were done on > 300 cells and from n = 3 technical replicates. Reported values represent means with SD. *p < 0.05 **p < 0.01 ***p < 0.001 by Welch’s *t* test. (**C**) SDS-PAGE of cell lysates from the time course represented in (A). All samples were standardized to the same OD600 prior to loading. The expected MWs of the three constructs indicated are: mCherry 26.8 kDa; mCh-McdB[wt] 44.6 kDa; mCh-McdB[-3] 44.5 kDa. Normalized intensities from the indicated bands were quantified from 3 biological replicates (*below*). Reported values represent means with SD. Data were analyzed via Welch’s *t* test.

### McdB[-3] causes mispositioned carboxysomes, likely due to an inability to interact with McdA

Having identified a mutant that solubilizes McdB condensates, without affecting structure or hexamerization, we set out to determine its influence on carboxysome positioning in *Se7942*. mNeonGreen (mNG) was N-terminally fused to either McdB[wt] or McdB[-3] and expressed at its native locus. The small subunit of Rubisco (RbcS) was C-terminally fused to mTurquoise (mTQ) to image carboxysomes. As shown previously (7), mNG-McdB[wt] supported well-distributed carboxysomes along the cell length (Fig. 8A, Fig. S9). The mNG-McdB[-3] strain, on the other hand, displayed carboxysome aggregates. However, it is important to note that McdA is a ParA/MinD family ATPase, which typically interact with their adaptor proteins via basic resides in the N-terminus of the adaptor protein, analogous to McdB (40, 41, 42, 43, 44, 45). Therefore, it is highly likely that one or more of the basic residues removed from McdB[-3] not only modulate condensate formation, but also mediate McdA interaction. A loss in McdA interaction would explain the carboxysome aggregation phenotype, as we have shown previously (7,12). To investigate this possibility, we knocked out McdA in the McdB[-3] mutant and found no significant differences in carboxysome mispositioning compared to the *ΔmcdA* strain alone (Fig. S9). Together, the data suggest that carboxysome aggregation in the McdB[-3] strain is due to this mutant’s inability to interact with McdA, and is not necessarily due to the effects these mutations have on McdB condensate formation.

**Figure 8:**
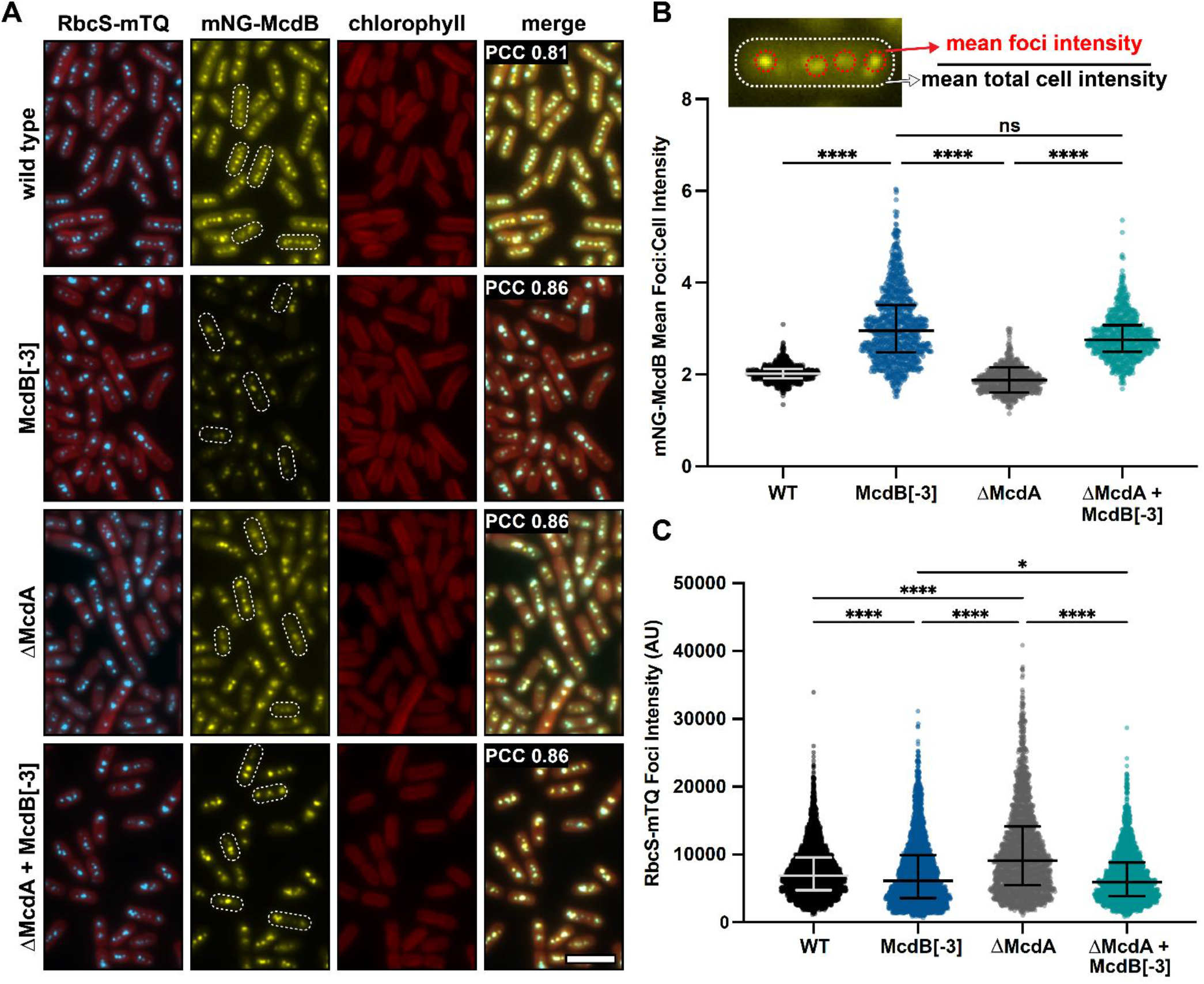
McdB[-3], which results in a high degree of condensate solubilization *in vitro* and in *E. coli*, alters the soluble fraction of McdB and carboxysome Rubisco levels *in vivo*. (**A**) Representative fluorescence microscopy images of the indicated strains. Scale bar = 5 µm and applies to all images. Pearson’s Correlation Coefficients (PCC) are shown for mNG-McdB and RbcS-mTQ for each strain. PCC values are means from > 10,000 cells over 10 fields of view. (**B**) Quantification of (mean foci intensity / mean total cell intensities) for mNG-McdB of n > 500 cells. Medians and interquartile ranges are displayed. **** p < 0.001 based on Kruskal-Wallis ANOVA. (**C**) Quantification of mean RbcS-mTQ foci intensity for n > 500 cells. Medians and interquartile ranges are displayed. *p < 0.05; ****p < 0.001 based on Kruskal-Wallis ANOVA.

### Condensate-defective McdB[-3] has a reduced cytoplasmic phase and associates with carboxysomes with lowered Rubisco content

Although carboxysome mispositioning by McdB[-3] cannot be directly ascribed to defects in McdB condensate formation specifically, two other observed phenotypes are not explained by a loss in McdA interaction. First, McdB[-3] still strongly colocalized with carboxysomes (PCC = 0.86 ± 0.01, n > 10,000 cells), similar to that of both McdB[wt] (PCC = 0.81 ± 0.01, n > 10,000 cells), and association did not change in the absence of McdA (PCC = 0.86 ± 0.02, n > 10,000 cells) (Fig. 8A). The data strongly suggests that condensate formation is not required for McdB to associate with carboxysomes.

Strikingly, however, the cytoplasmic phase observed for McdB[wt] was significantly lower for McdB[-3], both in the presence or absence of McdA (Fig. 8A). Indeed, when quantifying the intensity ratio of carboxysome-associated McdB to that of the whole cell, McdB[-3] showed a significant deviation in the ratio that was independent of McdA (Fig. 8B). The data show that while McdB condensate formation is not required for carboxysome association, without this activity, the cytoplasmic fraction of McdB notably declines.

Finally, we set out to directly determine the effect of McdB[-3] on the carboxysome itself by quantifying encapsulated Rubisco. Intriguingly, the McdB[-3] strains, with or without McdA, had carboxysomes with significantly lower Rubisco content as quantified by RbcS-mTQ intensity (Fig. 8C). This finding was particularly striking in the *ΔmcdA* background because, as we have shown previously, deletion of McdA results in increased RbcS-mTQ foci intensity due to carboxysome aggregation (46). But with McdB[-3], RbcS-mTQ intensity decreased, even with McdA deleted (Fig. 8C). Together, these data show that McdB[-3] increases the carboxysome-bound to soluble-McdB ratio and decreases Rubisco content in carboxysomes. These phenotypes are not explained by the loss of interaction with McdA, and are therefore potentially linked to defects in McdB condensate formation.

## DISCUSSION

In this report, we generate an initial structural model of *Se7942* McdB based on several empirical and predictive approaches. We define a tripartite domain architecture with an N-terminal IDR, a stable CC domain, and a CTD consisting of several α-helices (Fig. 1). We show that the CC dimerizes McdB and the CTD trimerizes the dimer, resulting in a trimer-of-dimers hexamer. The IDR had no impact on oligomerization (Fig. 2). Next, we found that McdB forms condensates via pH-dependent PSCP, where condensates show time-dependent viscoelastic properties and McdB forms pH-dependent clusters at concentrations far below the observed c_sat_ (Fig. 3). Using truncations, we found that the CC domain drives condensate formation, the IDR modulates solubility, and the CTD provides further valency. Therefore, all three domains are required for achieving wild-type levels of condensate formation (Fig. 4). We then identified positive residues in McdB important for stabilizing condensates. By performing scanning mutagenesis in both the IDR and CTD, we show that while mutations to the CTD destabilize the McdB hexamer, substituting out basic residues in the IDR solubilized condensates without affecting McdB structure or oligomerization (Fig. 5). These findings allowed us to design a series of mutants where the net charge of the IDR tuned McdB condensate solubility both *in vitro* (Fig. 6) and *in vivo* (Fig. 7). Lastly, we found that a solubilized McdB mutant, McdB[-3], impacts the carboxysome-bound to soluble McdB ratio in the cell, as well as Rubisco content in carboxysomes (Fig. 8). Overall, we determined McdB domain architecture, its oligomerization domains, regions required for condensate formation, and how to fine-tune condensate solubility, allowing us to link McdB condensate formation to potential functions *in vivo* (Fig. 9).

**Figure 9:**
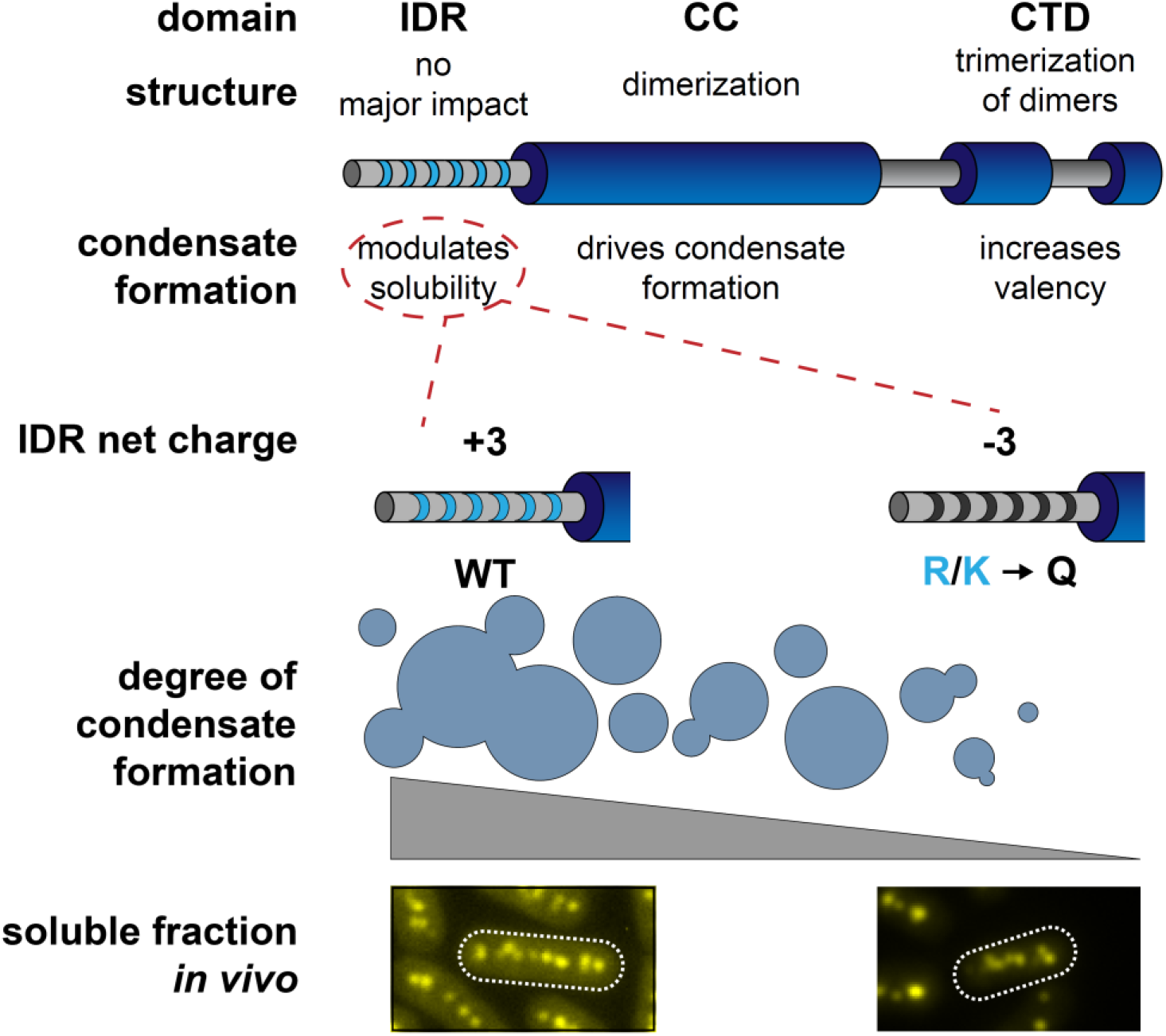
Proposed model of *Se7942* McdB domain structure and self-association. The central coiled-coil domain (CC) is necessary and sufficient for dimerization and driving condensate formation. The α-helical C-terminal domain (CTD) trimerize McdB dimers and increases the degree of condensate formation compared to the CC alone. The N-terminal IDR does not affect oligomerization and increases the degree of condensate formation compared to the CC alone. Substituting basic residues (K/R) in the IDR to glutamines (Q) can tune condensate solubility *in vitro* without affecting McdB oligomerization. These mutations allowed us to identify *in vivo* phenotypes correlated specifically to McdB phase separation, including the relative amount of soluble McdB.

### McdB condensate formation follows a nuanced, multi-domain mechanism

As the field of biomolecular condensates advances, more nuanced mechanisms are arising that describe combinations of condensate-driver domains, solubility modulators, and influences of oligomerization. We see here that different domains of McdB influence condensate formation via effects on solubility, self-association, and network formation. For many proteins, IDRs have been shown to be necessary and sufficient for driving condensate formation to a degree that is comparable to the full-length protein (47, 48, 49, 50, 51). The IDR of McdB, on the other hand, did not form condensates even at high protein concentrations and in the presence of a crowder. Furthermore, while deleting the IDR did not prevent McdB condensate formation, substituting only six basic residues in the IDR with glutamines solubilized condensates both *in vitro* and *in vivo*. In line with our findings, recent models have described how charged residues within IDRs can serve as key modulators of protein phase separation via mediating interactions with the solvent (52, 53, 54).

Glutamine-rich regions are known to be involved in stable protein-protein interactions such as in coiled-coils and amyloids (55, 56), and expansion of glutamine-rich regions in some proteins lead to amylogenesis and disease (57, 58). Thus, one potential caveat here is that introduction of glutamines may lead to amylogenesis for McdB. However, when we introduced glutamines into the IDR of McdB, solubility was increased both *in vitro* and *in vivo* without any impact on hexamerization. Our findings therefore underpin the importance of positive charge in the IDR specifically for stabilizing McdB condensates.

Oligomerization has also been found to influence protein condensate formation. For example, some proteins require oligomerization to provide the multivalency needed to form condensates (37, 59, 60), where some IDRs only induce condensate formation when fused to an oligomerizing domain (36). In some cases, oligomerization domains, like coiled-coils, drive condensate formation and are modulated by other domains (61), similar to our findings here with McdB. Such is the case for the bacterial protein PopZ, where an oligomerization domain forms condensates with solubility modulated by an IDR (19).

Truncated proteins have been useful in the study of biomolecular condensates. But it is important to note that using truncation data alone to dissect modes of condensate formation can lead to erroneous models since entire regions of the protein are missing. However, data from our truncation and substitution mutants were entirely congruent. For example, deletion of the CTD or substitutions to this region caused destabilization of the hexamer to a dimer, and deletion of the IDR or substitutions to this region caused solubilization of condensates without affecting hexamer formation. Furthermore, it should be noted that the McdB constructs used in our *in vitro* assays were free from fluorescent proteins, organic dyes, or other modifications that may influence phase separation. Therefore, the observed material properties of these condensates have full dependence on the McdB sequence.

### McdB homologs have polyampholytic properties between their N-and C-termini

In our previous bioinformatic study, we found that McdB homologs possess features that enable condensate formation, including intrinsic disorder, low hydrophobicity, biased amino acid compositions, and multivalency (8). Intriguingly, we also found that McdB proteins were potentially polyampholytes, with biphasic charge distributions between the NTD and CTD flanking the CC domain. For *Se7942* McdB, the N-terminal IDR has a pI of 10.8 and the CTD has a pI of 6.8, suggesting a potential electrostatic interaction between the two termini. The reason for such a shared feature was not obvious, but we proposed this polyampholytic nature was important for McdB self-association. A structure of the CC domain of a plasmid-encoded McdB-like protein from the cyanobacterium *Cyanothece sp.* PCC 7424 displayed an antiparallel association to form a dimer (62). Antiparallel dimerization of the CC domains of *Se7942* McdB would align these oppositely charged extensions. Consistently, our truncation data provides evidence suggesting a condensate-stabilizing interaction between the N-terminal IDR and the CTD. Condensates from the IDR+CC construct, which lack the CTD, were highly dynamic. But with the CTD present, condensates fused slowly. We were unable to dissect how the CTD contributes to condensate formation as substitutions to the CTD caused hexamer destabilization. However, these results have set the stage for several future studies that will (*i*) probe the orientation of McdB subunits within the hexamer, (*ii*) determine how basic residues in the IDR influence McdB condensate formation at a molecular level, and (*iii*) identify residues in the CTD that may influence condensate formation.

### Partner Proteins of ParA/MinD ATPases form condensates with functional implications

McdA is a member of the ParA/MinD family of ATPases that position a wide array of cellular cargos, including plasmids, chromosomes, and the divisome (10, 11, 12). A partner protein acts as the adaptor, linking the positioning ATPase to its respective cargo. McdB is the adaptor protein that links McdA to the carboxysome. Here we show that McdB forms condensates both *in vitro* and *in vivo*, and the solubility of these condensates influences the nature of McdB association with carboxysomes. Intriguingly, condensate formation by other adaptor proteins has been proposed to play important roles in mediating interactions with other cargos positioned by ParA/MinD ATPases. For example, ParA ATPases are responsible for the spatial regulation of chromosomes and plasmids that are bound by the adaptor protein ParB (10, 11, 12). The exact nature of the interaction between ParB and DNA remains a vibrant area of research (63), but recent reports show that ParB-DNA complexes behave as dynamic, liquid-like condensates both *in vitro* (64) and *in vivo* (65), implementing condensate formation as an underlying mechanism. Another example is the co-complex of adaptor proteins, PomX and PomY, for the ParA/MinD ATPase called PomZ. PomXYZ is responsible for spatially regulating division sites in some bacteria (66). A recent study has shown that PomY forms condensates that nucleate GTP-dependent FtsZ polymerization, suggesting a novel mechanism for positioning cell division (67). It is intriguing to speculate why such disparate adaptor proteins could use condensate formation as an underlying mechanism in localizing to vastly different cargos. The ability to undergo changes in density at specific locations and times within the cell may confer an advantage to these highly dynamic spatial regulation systems.

### pH as a potential underlying regulator for McdB condensate solubility and its association with carboxysomes

In carbon-fixing organisms, the collection of processes that contribute to efficient carbon fixation are referred to as the <u>c</u>arbon <u>c</u>oncentrating <u>m</u>echanism (CCM). The development of a model for the cyanobacterial CCM has provided insight into how different features, such the presence of carboxysomes, affect overall carbon capture (68). It was shown that incorporation of a pH flux into CCM models provides values more consistent with experimentation (69). This updated ‘pH-aware’ model suggests that the carboxysome lumen maintains a lower pH than the cytoplasm via Rubisco proton production; with the cytoplasm being ∼ pH 8.5 and the carboxysome lumen being ∼ pH 7.5 (69, 70).

Here, we show pH as a major regulator of McdB condensate solubility *in vitro*. Furthermore, we report a potential link between McdB condensate solubility and regulating both the carboxysome-bound and soluble fractions of McdB as well as Rubisco content within carboxysomes. We speculate that intracellular differences in pH may influence McdB localization and function by modulating self-associating interactions that underlie McdB condensate formation. In this model, condensation-deficient McdB[-3] associates with carboxysomes, but no longer dynamically exchanges with the cytoplasmic fraction. Stable McdB[-3] association with carboxysomes may influence the dynamic process of carboxysome assembly and maintenance, which would explain the decreased Rubsico content observed here.

Future studies will determine the nature of McdB association with carboxysomes, and how condensate formation influences this association. For example, we have shown that McdB strongly associates with carboxysome shell proteins via bacterial two-hybrid assays (7). It is attractive to speculate that McdB undergoes pre-wetting interactions with the 2D surface of the carboxysome shell, which then nucleates surface-assisted condensation. Such 2D interactions would significantly impact McdB phase boundaries (71). Teasing apart the stable protein-protein interactions between McdB and carboxysomes from the dynamic processes governing condensate solubility will therefore be of significant importance.

### Tunable protein condensates as useful tools for synthetic biology

A useful property of biomolecular condensates is the ability to regulate enzyme activity (72). Specific chemistries within condensates can affect the degree to which certain metabolites and enzymes are soluble within the dense phase. Thus, condensates can serve as reaction centers that regulate the overall metabolism of a cell by transiently altering the activities of key reactions. For example, it has been shown that certain scaffolding proteins can form phase separated condensates with Rubisco (73, 74). It is speculated that these Rubisco condensates were the original CCM, which then led to the evolution of carboxysomes and the modern CCM (70). An exciting future direction for the field of biomolecular condensates is the prospect of designing condensate forming scaffolding proteins that can recruit specific enzymes, such as Rubisco, and implementing these designer enzyme-condensates in synthetic cells to engineer metabolism (19, 20, 72).

Here, we find that the IDR of McdB, which does not itself drive condensate formation, is amenable to mutations that fine-tune McdB condensate properties. The bacterial protein PopZ has already been engineered to fine-tune condensate formation and has also been developed as a tool called the “PopTag”, which endows condensate forming activity to fusion proteins expressed in a variety of cell types, including human cells (20). Expanding the repertoire of protein condensates with tunable properties will further advance our potential for designing and engineering reaction isolation and cellular metabolisms.

## ACKNOWLEDGEMENTS

We would like to thank Dr. JK Nandakumar and Ritvija Agrawal for training and allowing us to use their SEC-MALS system. Dr. Henriette Remmer at the University of Michigan Proteomics & Peptide Synthesis Core for help with N-terminal sequencing. The National Crystallization Center at the Hauptman-Woodward Medical Research Institute for performing crystallization buffer screens. This work was supported by the National Science Foundation to A.G.V. (NSF CAREER Award No. 1941966), Rackham Graduate Student Research Grant to J.L.B., and from research initiation funds provided by the MCDB Department to A.G.V.

## MATERIALS AVAILABILITY STATEMENT

Any materials used in this study are described in detail in Experimental Procedures or can be accessed by contacting the authors.

## DATA AVAILABILITY STATEMENT

The data that supports the findings of this study are available in the supplementary material of this article or can be accessed by contacting the authors.

## EXPERIMENTAL PROCEDURES

### Protein expression and purification

Wild-type and mutant variants of McdB were expressed with an N-terminal His-SUMO tag off a pET11b vector in *E. coli* BL21-AI (Invitrogen). All cells were grown in LB + carbenicillin (100 µg/mL) at 37°C unless otherwise stated. One liter cultures used for expression were inoculated using overnight cultures at a 1:100 dilution. Cultures were grown to an OD_600_ of 0.5 and expression was induced using final concentrations of IPTG at 1 mM and L-arabinose at 0.2%. Cultures were grown for an additional 4 hours, pelleted, and stored at -80°C.

Pellets were resuspended in 30 mL lysis buffer [300 mM KCl; 50 mM Tris-HCl pH 8.4; 5 mM BME; 50 mg lysozyme (Thermo-Fischer); protease inhibitor tablet (Thermo-Fischer)] and sonicated with cycles of 10 seconds on, 20 seconds off at 50% power for 7 minutes. Lysates were clarified via centrifugation at 15,000 rcf for 30 minutes. Clarified lysates were passed through a 0.45 µm filter and loaded onto a 1 mL HisTrap HP (Cytiva) equilibrated in buffer A [300 mM KCl; 50 mM Tris-HCl pH 8.4; 5 mM BME]. Columns were washed with 5 column volumes of 5% buffer B [300 mM KCl; 20 mM Tris-HCl pH 8.4; 5 mM BME; 500 mM imidazole]. Elution was performed using a 5-100% gradient of buffer B via an AKTA Pure system (Cytiva). Peak fractions were pooled and diluted with buffer A to a final imidazole concentration of < 100 mM. Ulp1 protease was added to a final concentration of 1:100 protease:sample, and incubated overnight at 23°C with gentle rocking. The pH was then adjusted to ∼10 and samples were concentrated to a volume of < 5 mL, passed through a 0.45 µm filter and passed over a sizing column (HiLoad 16/600 Superdex 200 pg; Cytiva) equilibrated in buffer C [150 mM KCl; 20 mM CAPS pH 10.2; 5 mM BME; 10% glycerol]. Peak fractions were pooled, concentrated, and stored at -80°C.

### Proteolysis and N-terminal sequencing

Proteolysis was performed on *Se7942* McdB at 30 µM in buffer containing 150 mM KCl, 50 mM HEPES pH 7.7, and 2 mM BME. Trypsin protease (Thermo-Fischer) was added at a 1:100 ratio of protease:protein. The reaction was incubated at 30°C and samples were quenched at the indicated time points by diluting into 4X Laemmli SDS-PAGE sample buffer containing 8% SDS. Degradation over time was visualized by running time points on a 4–12% Bis-Tris NuPAGE gel (Invitrogen) and staining with InstantBlue Coomassie Stain (Abcam).

Bands that were N-terminally sequenced were separated via SDS-PAGE as above, but transferred to a PVDF membrane (Bio-Rad) prior to staining. Transfer of bands was performed using a Trans-Blot Turbo Transfer System (Bio-Rad). N-terminal sequences of these bands were then determined using Edman degradation.

### Circular dichroism

For all protein samples analyzed, far-UV CD spectra were obtained using a J-1500 CD spectrometer (Jasco). All measurements were taken with 250 µL of protein at 0.25 mg/mL in 20 mM KPi, pH 8.0. Measurements were taken using a quartz cell with a path length of 0.1 cm. The spectra were acquired from 260 to 190 nm with a 0.1 nm interval, 50 nm/min scan speed, and at 25°C unless otherwise stated.

### Microscopy of protein condensates

Samples for imaging were set up in 16 well CultureWells (Grace BioLabs). Wells were passivated by overnight incubation in 5% (w/v) Pluronic acid (Thermo-Fischer), and washed thoroughly with the corresponding buffer prior to use. All condensate samples were incubated for 30 minutes prior to imaging unless otherwise stated. For experiments where samples were imaged across pH titrations, the following buffers were used: phosphate buffer for pH 6.3-6.7, HEPES for pH 7.2-7.7, and Tris-HCl for 8.2-8.6. Imaging of condensates was performed using a Nikon Ti2-E motorized inverted microscope (60 × DIC objective and DIC analyzer cube) controlled by NIS Elements software with a Transmitted LED Lamp house and a Photometrics Prime 95B Back-illuminated sCMOS Camera. Image analysis was performed using Fiji v 1.0.

### Fluorescence recovery after photobleaching (FRAP)

All FRAP measurements were performed using the indicated protein concentration with the addition of 1:1000 mNG-McdB based on molarity. All fluorescence imaging was performed using a Nikon Ti2-E motorized inverted microscope controlled by NIS Elements software with a SOLA 365 LED light source, a 100× objective lens (Oil CFI60 Plan Apochromat Lambda Series for DIC), and a Photometrics Prime 95B Back-illuminated sCMOS camera. mNG signal was acquired using a “GFP” filter set [excitation, 470/40 nm (450 to 490 nm); emission, 525/50 nm (500 to 550 nm); dichroic mirror, 495 nm]. Bleaching was conducted with a 405-nm laser at 40% power (20 mW) with a 200-μs dwell time. Recovery was monitored with a time-lapse video with 5 sec intervals for 20 mins. Image analysis was done in Fiji v 1.0. Intensities from bleached regions of interest (ROIs) were background subtracted and normalized using an unbleached condensate to account for any full field of view photobleaching. The values for each condensate were then normalized such that a value of 1 was set to the pre-bleach intensity and a value of 0 was set to the intensity immediately post-bleaching. Data were exported, further tabulated, graphed, and analyzed using GraphPad Prism 9.0.1 for macOS (GraphPad Software, San Diego, CA, www.graphpad.com).

### Dynamic light scattering (DLS)

All sizing and polydispersity measurements were carried out on an Uncle by Unchained Labs (USA) at 25°C in triplicate. Samples were adjusted to the indicated concentrations in 150 mM KCl and 20 mM of the following buffers based on pH: HEPES, pH 7.2; Tris-HCl, pH 8.2; CAPS, pH 10.2. Samples were analyzed both before and after a centrifugation step at 20,000 rcf for 10 min as indicated. Data were exported, further tabulated, graphed, and analyzed using GraphPad Prism 9.0.1 for macOS (GraphPad Software, San Diego, CA, www.graphpad.com).

### Size-exclusion chromatography (SEC)

SEC was performed on full-length and truncated McdB proteins using a Superdex 200 Increase 10/300 GL (Cytiva) column connected to an AKTA pure system (Cytiva). 500 µL of sample at 1.5 mg/mL protein was passed through the column at 0.4 mL/min in buffer [150 mM KCl; 20 mM Tris-HCl pH 8.2] while monitoring absorbance at 220 nm.

### SEC coupled to multi-angled light scattering (SEC-MALS)

For each sample analyzed, 50 µL at 1.5 mg/ml was passed over an SEC column (PROTEIN KW-804; Shodex) at a flow rate of 0.4 mL/min in buffer containing 150 mM KCl and 20 mM Tris-HCl, pH 8.2. Following SEC, the samples were analyzed using an A280 UV detector (AKTA pure; Cytiva), the DAWN HELEOS-II MALS detector with an internal QELs (Wyatt Technology), and the Optilab T-rEX refractive index detector (Wyatt Technology). The data were analyzed to calculate mass using ASTRA 6 software (Wyatt Technology). Bovine serum albumin was used as the standard for calibration.

### Phase diagrams

Data for phase diagrams was collected using an Infinite M200 PRO plate reader (Tecan). Samples were set up in 96 well glass-bottom plates (Thomas Scientific) and absorbance at 350 nm was measured as previously described (15). Reported values are averages of triplicates with buffer blanks subtracted, and error bars representing standard deviations. Protein concentration, KCl concentration, and pH values varied as indicated, but for each pH value tested, 20 mM of the following buffers were used: phosphate buffer for pH 6.3-6.7, HEPES for pH 7.2-7.7, and Tris-HCl for pH 8.2-8.6.

### Quantification of phase separation via centrifugation

Centrifugation was used to quantify the degree to which McdB and its variants condensed under certain conditions, as previously described (15). Briefly, 100 µL of sample was incubated at the conditions specified for 30 minutes, and then centrifuged at 16,000 rcf for 2 minutes. The supernatant was removed and the pellet resuspended in an equal volume of McdB solubilization buffer [300 mM KCl, 20 mM CAPS pH 10.2]; McdB does not condense at pH 10.2. Samples were then diluted into 4X Laemmli SDS-PAGE sample buffer. Pellet and supernatant fractions were visualized on a 4–12% Bis-Tris NuPAGE gel (Invitrogen) by staining with InstantBlue Coomassie Stain (Abcam) for 1 hour and then destaining in water for 14-16 hours. The intensities of the bands were quantified using Fiji v 1.0 and resultant data graphed using GraphPad Prism 9.0.1 for macOS (GraphPad Software, San Diego, CA, www.graphpad.com).

### Expression and visualization of mCherry fusions in E. coli

All constructs were expressed off a plasmid from a pTrc promoter in *E. coli* MG1655. Overnight cultures grown in LB + carbenicillin (100 µg/mL) were diluted at 1:100 into AB medium + carbenicillin (100 µg/mL) supplemented with (0.2% glycerol; 10 µg/mL thiamine; 0.2% casein; 25 µg/mL uracil). Cultures were grown at 37°C to an OD600 = 0.3 and induced with 1 mM IPTG. Following induction, cultures were grown at 37°C and samples taken at the indicated time points.

Cells used for imaging were prepared by spotting 3 µL of cells onto a 2% UltraPure agarose + AB medium pad on a Mantek dish. Images were taken using Nikon Ti2-E motorized inverted microscope controlled by NIS Elements software with a SOLA 365 LED light source, a 100X Objective lens (Oil CFI Plan Apochromat DM Lambda Series for Phase Contrast), and a Hamamatsu Orca Flash 4.0 LT + sCMOS camera. mCherry signal was imaged using a “TexasRed” filter set (C-FL Texas Red, Hard Coat, High Signal-to-Noise, Zero Shift, Excitation: 560/40 nm [540-580 nm], Emission: 630/75 nm [593-668 nm], Dichroic Mirror: 585 nm). Image analysis was performed using Fiji v 1.0.

To monitor expression levels, cells were harvested via centrifugation at the indicated time points, and resuspended in 4X Laemmli SDS-PAGE sample buffer to give a final OD600 = 4. Samples were boiled at 95°C and 10 µL were then run on a 4–12% Bis-Tris NuPAGE gel (Invitrogen). Bands were visualized by staining with InstantBlue Coomassie Stain (Abcam) for 1 hour and then destaining in water for 14-16 hours. Quantifying the normalized band intensities was performed using Fiji v 1.0.

### Growth and transformation of S. elongatus PCC 7942

All *S. elongatus* (ATCC 33912) strains were grown in BG-11 medium (Sigma) buffered with 1 g/L HEPES, pH 8.3. Cells were incubated with the following growth conditions: 60 μmol m–2 s–1 continuous LED 5600 K light, 32°C, 2% CO2, and shaking at 130 RPM. Transformations of *S. elongatus* cells were performed as previously described (75). Transformants were plated on BG-11 agar with 12.5 µg/ml kanamycin. Single colonies were picked and transferred liquid BG-11 medium with corresponding antibiotic concentrations. Complete gene insertions and absence of the wild-type gene were verified via PCR, and cultures were removed from antibiotic selection prior to imaging.

### Live cell fluorescence microscopy and analysis

100 µL of exponentially growing cells (OD750 ∼ 0.7) were harvested and spun down at 4000 rcf for 1 min and resuspended in 10 µl fresh BG-11. 2 µl of the resuspension were then spotted on 1.5% UltraPure agarose (Invitrogen) + BG-11 pad on a 35-mm glass-bottom dish (MatTek Life Sciences). All fluorescence and phase-contrast imaging were performed using a Nikon Ti2-E motorized inverted microscope controlled by NIS Elements software with a SOLA 365 LED light source, a 100× objective lens (Oil CFI Plan Apochromat DM Lambda Series for Phase Contrast), and a Photometrics Prime 95B back-illuminated sCMOS camera or Hamamatsu Orca-Flash 4.0 LTS camera. mNG-McdB variants were imaged using a “YFP” filter set (C-FL YFP, Hard Coat, High Signal-to-Noise, Zero Shift, Excitation: 500/20 nm [490–510 nm], Emission: 535/30 nm [520–550 nm], Dichroic Mirror: 515 nm). RbcS-mTQ-labeled carboxysomes were imaged using a “CFP” filter set (C-FL CFP, Hard Coat, High Signal-to-Noise, Zero Shift, Excitation: 436/20 nm [426–446 nm], Emission: 480/40 nm [460-500 nm], Dichroic Mirror: 455 nm). Chlorophyll was imaged using a “TexasRed” filter set (C-FL Texas Red, Hard Coat, 583 High Signal-to-Noise, Zero Shift, Excitation: 560/40 nm [540-580 nm], Emission: 630/75 nm 584 [593-668 nm], Dichroic Mirror: 585 nm).

Image analysis including cell segmentation, quantification of foci number, intensities, and spacing were performed using Fiji plugin MicrobeJ 5.13n (8) (76). Cell perimeter detection and segmentation were done using the rod-shaped descriptor with default threshold settings. Carboxysome foci detection was performed using the point function with tolerance of 700 and the sharpen image filter selected. McdB foci detection was performed using the smoothed foci function with tolerance of 100, Z-score of 3, and the minimum image filter selected. Pearson’s correlation coefficients were calculated using ImageJ plugin JaCoP (77), and reported values represent means and standard deviations from > 10,000 cells over 10 fields of view. Data were exported, further tabulated, graphed, and analyzed using GraphPad Prism 9.0.1 for macOS (GraphPad Software, San Diego, CA, www.graphpad.com).

## SUPPLEMENTAL FIGURES

**Figure S1:**
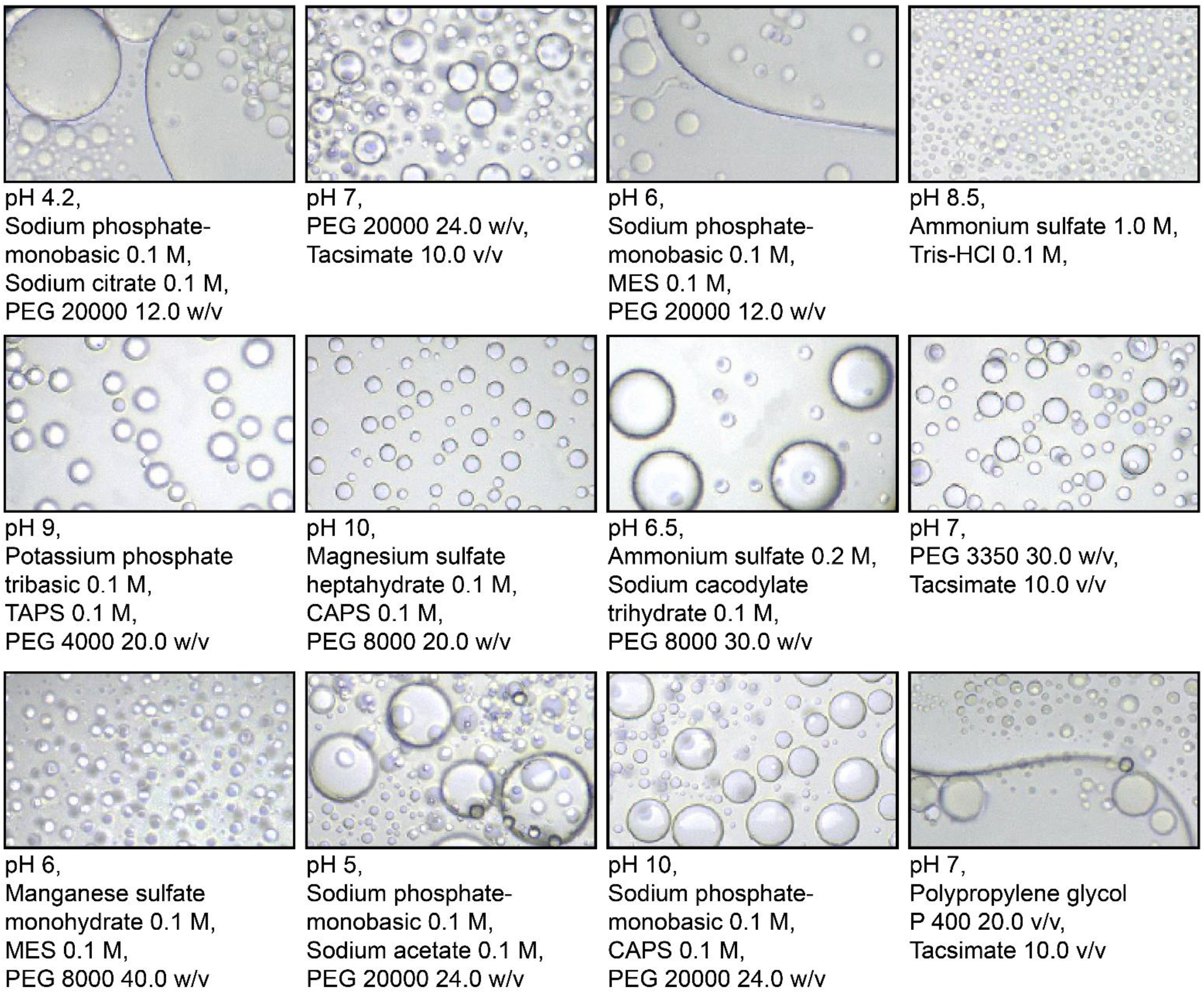
Phase separation of *Se7942* McdB across a range of buffer conditions during crystal screens. Images taken during buffer screens for crystallography. McdB at 10 mg/mL in (50 mM KCl; 10 mM CAPS pH 10.2) was diluted into the buffers indicated below each image. All images shown are at the same final concentration and magnification. Images were taken after 24-36 hours post dilution.

**Figure S2:**
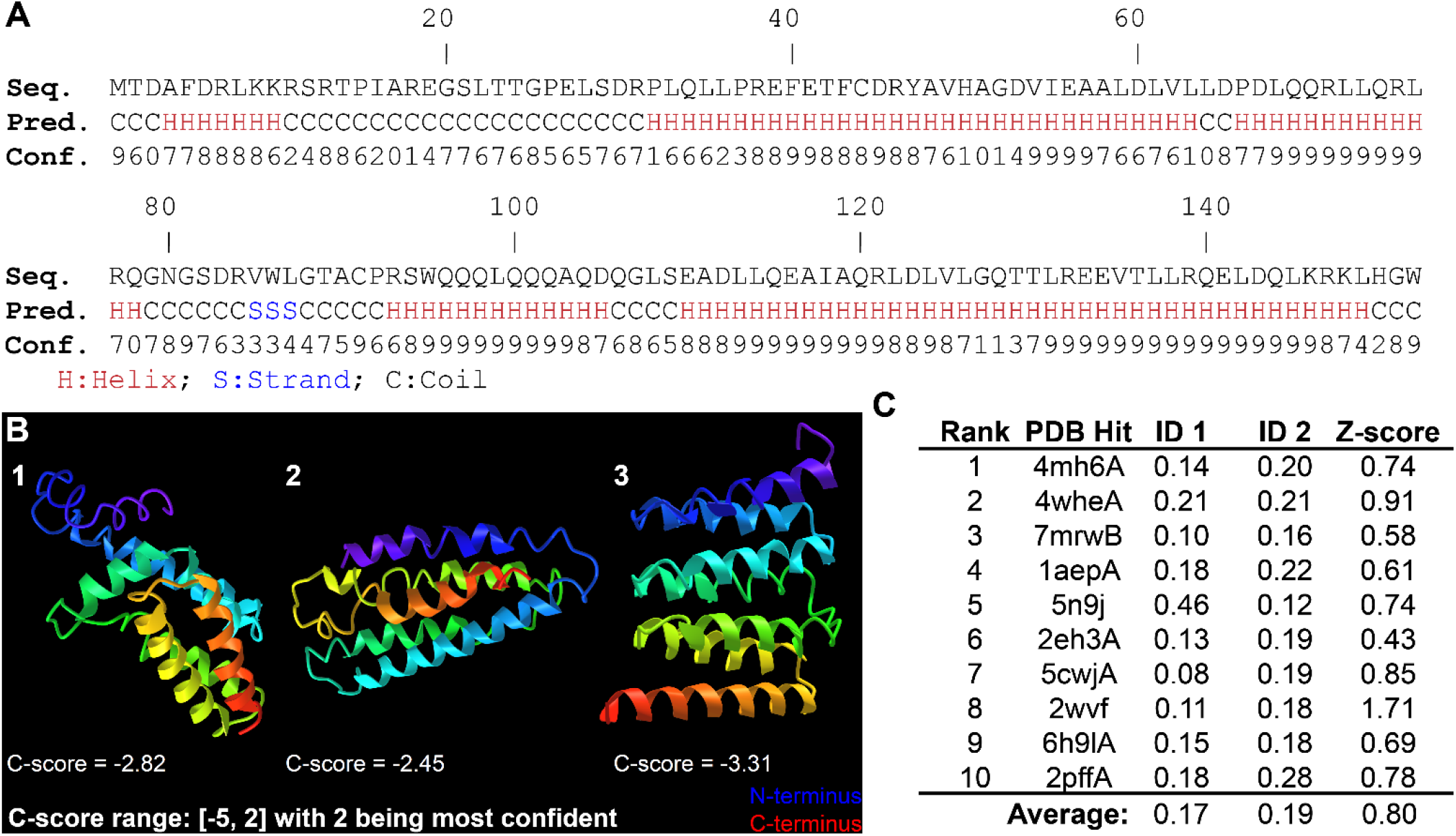
I-TASSER predictions for *Se7942* McdB. (**A**) McdB amino acid sequence and associated secondary structure predictions by I-TASSER. Each residue has a confidence score that ranges from 0 (least confident) to 9 (most confident). (**B**) Top three final models generated by I-TASSER. Each model is given a C-score that ranges from [-5, 2] with -5 being the least confident and 2 being the most. N-termini are colored blue and C-termini are colored red. (**C**) A table listing the top 10 PDB templates identified by I-TASSER, which were used for generating the models in panel B. ID1 is the percent sequence identity of the templates in the threading-aligned region with the query sequence. ID2 is the percent sequence identity of the whole template chains with the query sequence. Z-scores are a normalized score of the threading alignments. Alignments with a Z-score > 1 equates to good alignment. Overall, I-TASSER was unsuccessful in predicting a structure for McdB.

**Figure S3:**
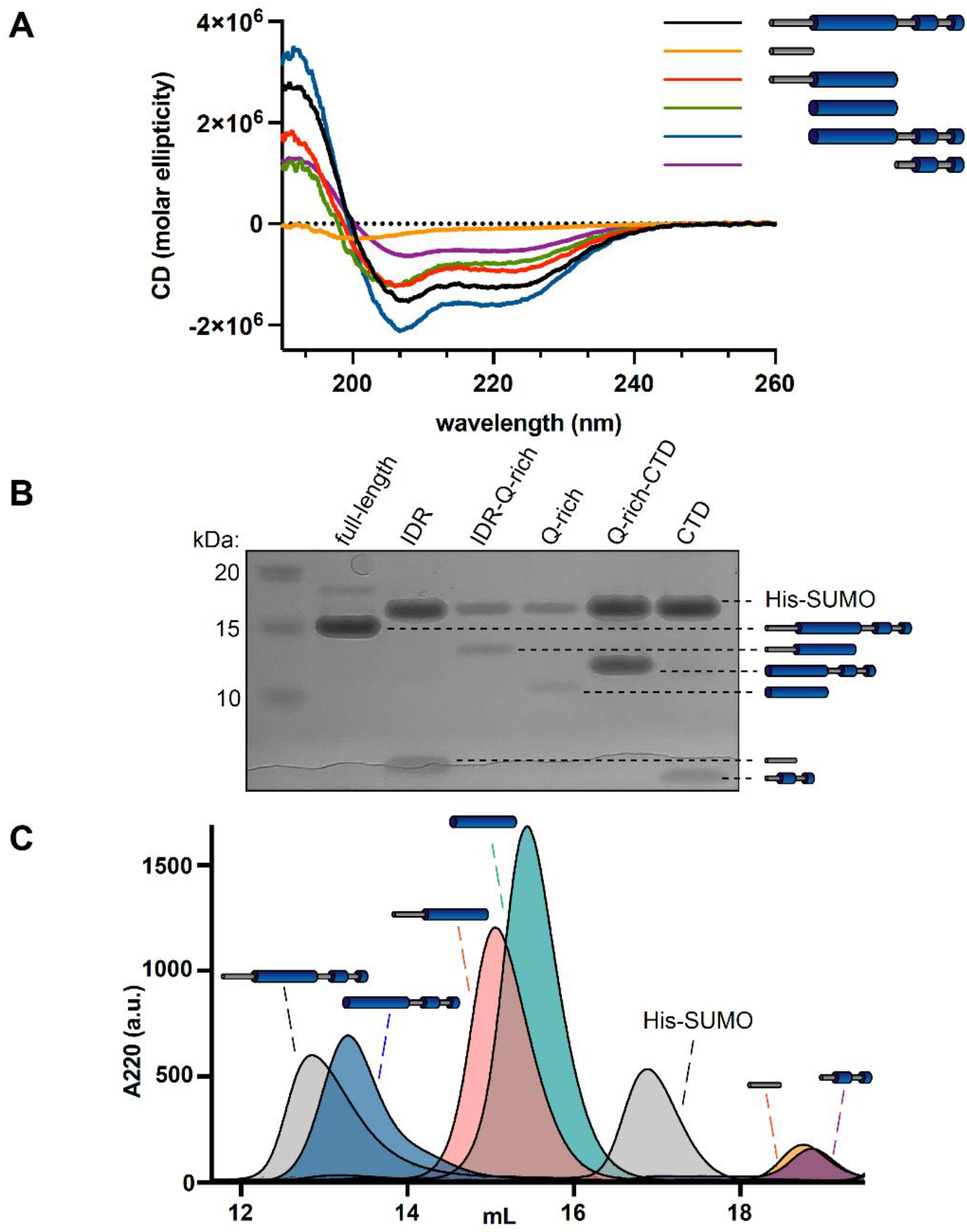
McdB truncations have unique secondary structures and display different oligomeric states. (**A**) CD spectra of full-length McdB and truncations. Curves from Fig. 2B are overlayed onto a single graph. (**B**) SDS-PAGE analysis shows that full-length McdB and all truncation mutants run at a lower molecular weight compared to the His-SUMO solubility tag. (**C**) Size exclusion chromatography (SEC) showed that full-length McdB and the CC+CTD domain have similar elution profiles, suggesting similar oligomeric forms. The CC domain with and without IDR also elute similarly but after full-length and before His-SUMO, suggesting an intermediate oligomer. The IDR and CTD mutants eluted after the His-SUMO tag, showing they remain monomeric.

**Figure S4:**
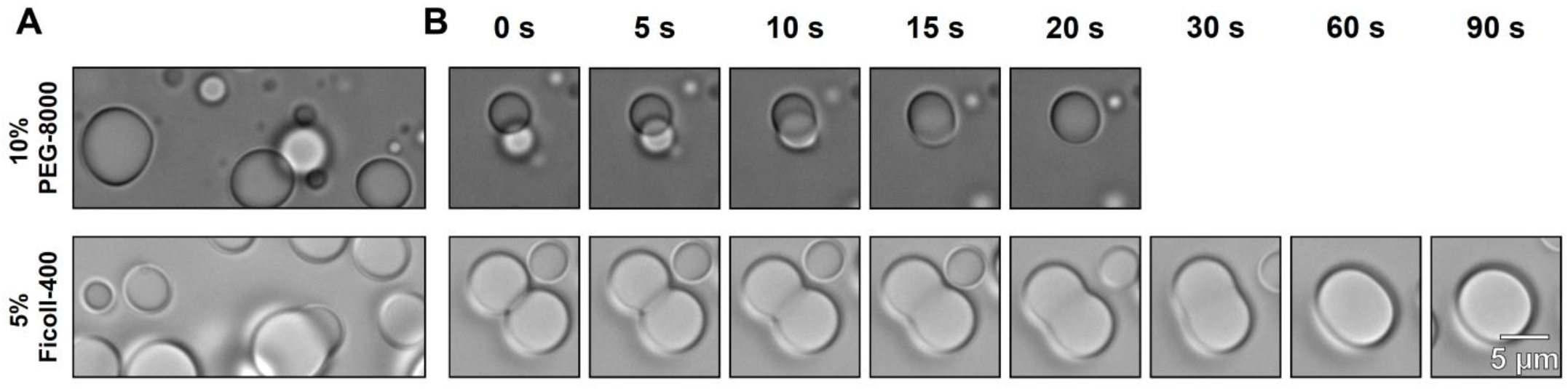
McdB forms liquid-like condensates in both Ficoll and PEG. (**A**) Representative DIC microscopy images for WT McdB at 100 µM in 100 mM KCl, 20 mM HEPES pH 7.2, and the addition of the indicated crowding agent. (**B**) Time course of images from (A) show that condensates fuse and relax into spheres on similar timescales, regardless of the crowding agent used.

**Figure S5:**
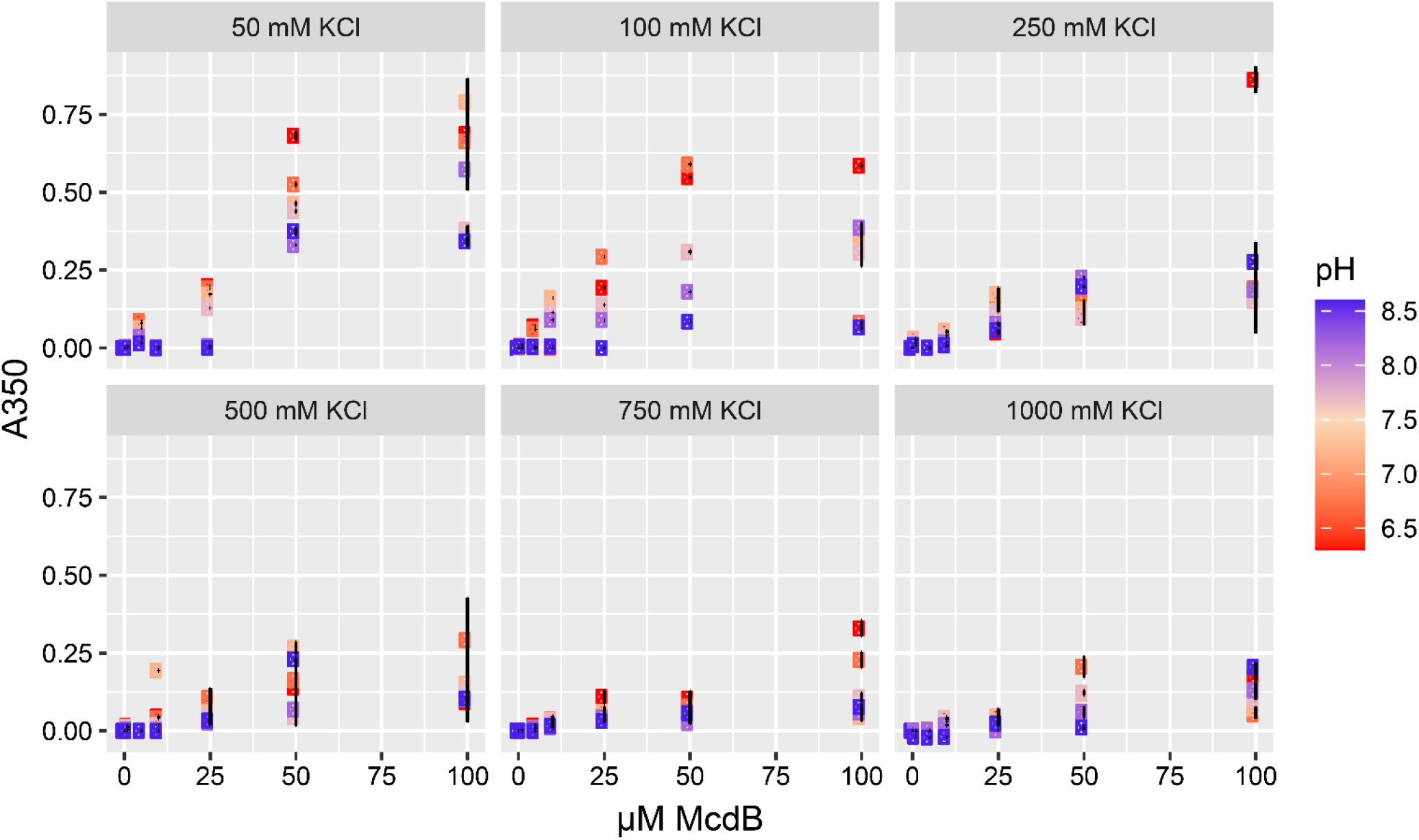
Multi-dimensional phase diagrams for *Se7942* McdB. Turbidity-based phase diagrams for McdB across varying protein concentration, KCl concentration, and pH. Data points represent the mean and error bars represent SD from at least three technical replicates. Turbidity monitored at A = 350 nm.

**Figure S6:**
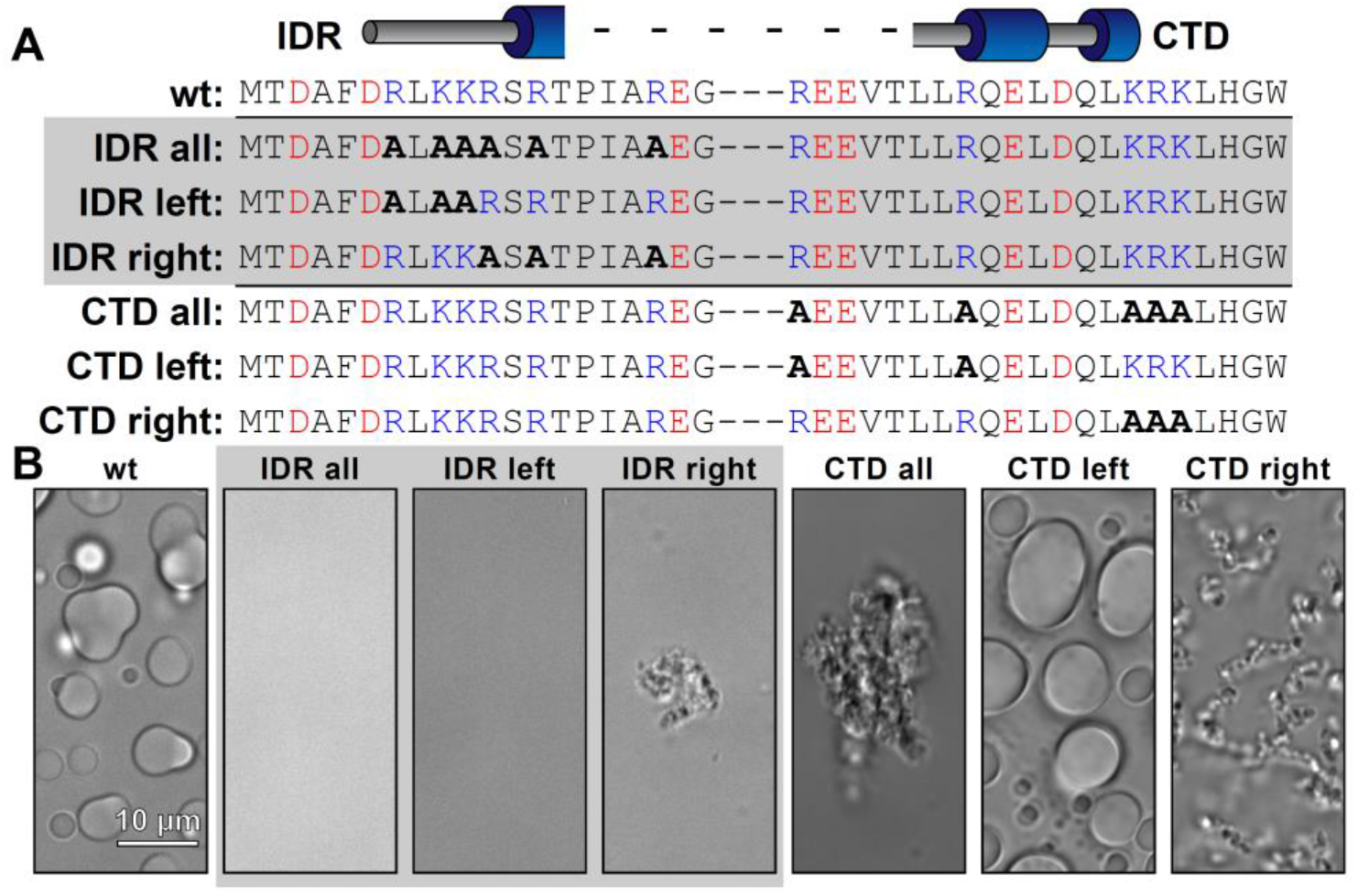
Alanine-scanning of basic residues in the N- and C-termini of McdB. (**A**) Table showing the sequence of WT McdB compared to the terminal A-substitution mutants. Acidic- and basic-residues are colored red and blue, respectively. A-substitutions are bolded. (**B**) Representative DIC microscopy images of all constructs listed in (A). Scale bar applies to all images.

**Figure S7:**
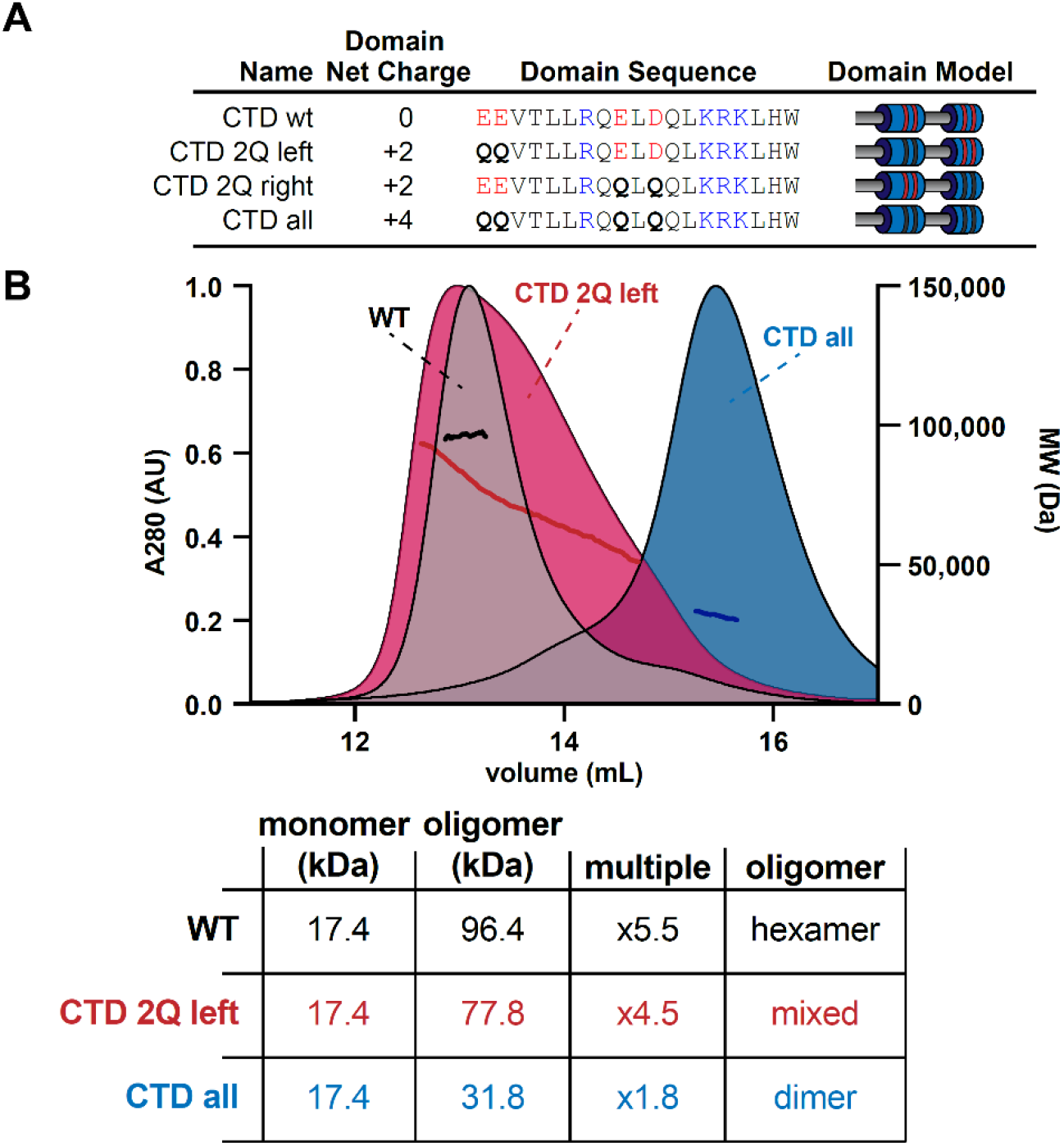
Mutations to the CTD destabilize the trimer-of-dimers hexamer. (**A**) Table showing the net charge and amino acid sequence of wild-type McdB compared to the Q substitution mutants in the CTD. Acidic and basic residues are colored red and blue, respectively. Q-substitutions are bolded. Graphical models of the McdB variants are also provided. (**B**) SEC-MALS graphs of the indicated variants (*top*) with a table summarizing results (*below*). Note the CTD 2Q left MALS data spans the MW from hexamer range to dimer range. The CTD 2Q right variant formed insoluble aggregates and is not shown.

**Figure S8:**
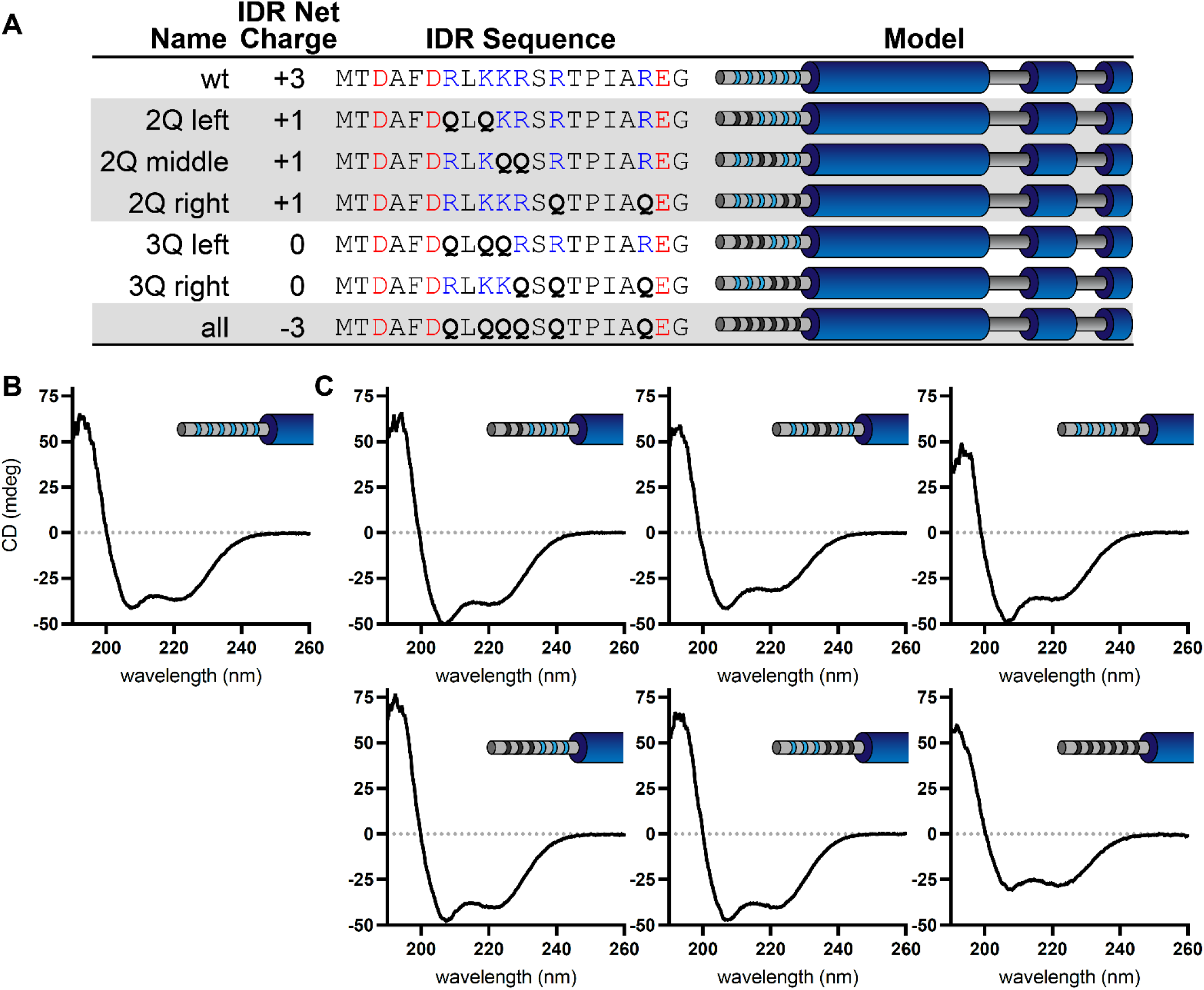
CD spectra of wild-type McdB and N-terminal glutamine substitution mutants. (**A**) Table showing the net charge and N-terminal IDR sequence of wild-type McdB compared to the glutamine-substitution mutants. Acidic- and basic-residues in the IDR are colored red and blue, respectively. Glutamine-substitutions are bolded. Graphical models of the McdB variants are also provided where blue stripes represent the six basic residues in the IDR. Black stripes represent the location of the glutamine substitutions. CD spectra of both (**B**) wild-type McdB and (**C**) mutants with the indicated glutamine substitutions in the N-terminal IDR of McdB.

**Figure S9:**
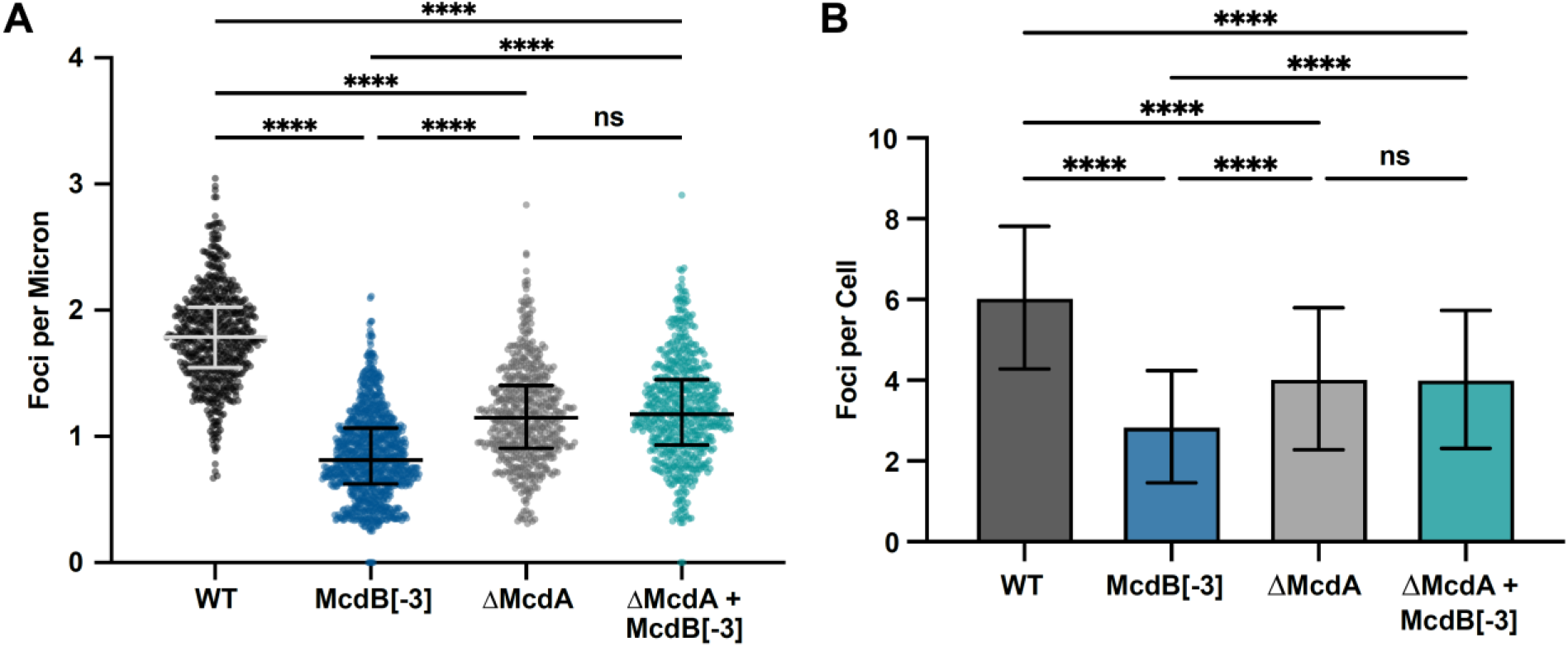
Deletion of McdA causes no additional loss of carboxysome positioning in McdB[-3] strain. (**A**) Quantification of RbcS-mTQ foci per micron from n > 500 cells. Medians and interquartile ranges are displayed. **** p < 0.001 based on Kruskal-Wallis ANOVA. (**B**) Quantification of RbcS-mTQ foci per cell for n > 500 cells. Medians and interquartile ranges are displayed. ****p < 0.001 based on Kruskal-Wallis ANOVA.

